# Metalink: Joint taxonomic and functional profiling of metagenomes and metatranscriptomes across resolution scales

**DOI:** 10.64898/2026.10.13.759005

**Authors:** Liuyang Li, Yinzhao Wang

## Abstract

Metagenomic and metatranscriptomic analyses remain largely fragmented: taxonomic and functional profiling are performed separately, often at a single resolution—reads, contigs, or MAGs—yielding separate species or function lists rather than mechanistic insights. Here, we developed Metalink, a tool that unifies taxonomic and functional profiling into coupled outputs across all three resolution levels. Metalink is built on three innovations. First, we constructed a Last Common Function-based functional Hierarchy (LCFH) from ∼424 million proteins, organizing KEGG Orthologs into a tree structure that enables LCA-like functional assignment. Second, a unified framework integrates the taxonomy of Genome Taxonomy Database (GTDB), the International Committee on Taxonomy of Viruses (ICTV), and NCBI for consistent classification across reads, contigs, and MAGs. Third, a multi-level integration strategy with overlap-based deduplication balances accuracy and completeness. At the read level, Metalink exceeds both HUMAnN3 and eggNOG-Mapper in functional assignment accuracy while running ∼30-fold faster and 1,000-fold faster, respectively; at the contig level, its functional profiles remain highly consistent with eggNOG-Mapper. Multi-level integration further improves taxonomic resolution over single-level analyses and outperforms leading taxonomic profilers on the CAMI II benchmarks. Metalink’s joint outputs move the analysis of metagenomes and metatranscriptomes from cataloging species and functions toward mechanistic inference of species–function–environment triadic associations. By enabling these integrated analyses across resolution scales, Metalink provides a general framework for linking community composition to its functional potential across diverse ecosystems.

## Introduction

Microbial communities drive essential biogeochemical processes across virtually all ecosystems on Earth, yet the vast majority of these organisms remain uncultivated and poorly characterized. Metagenomic and metatranscriptomic (meta’omic) sequencing have emerged as indispensable tools for accessing both the composition and functional potential of these communities without prior cultivation^1,2^. Despite the wealth of data generated by these approaches, understanding how community composition relates to function remains a major challenge in microbial ecology^3^. Most existing analytical tools address either taxonomic profiling or functional profiling in isolation^4–6^. Taxonomic profiling methods include marker-based approaches such as MetaPhlAn^1^ and mOTUs^7^, *k*-mer-based approaches such as Kraken2^8^, ganon^9^, sylph^10^, Centrifuge^11^ and Kaiju^12^, which infer community composition using diverse reference indexing strategies, whereas functional profiling and annotation methods such as HUMAnN^2^, MG-RAST^13^, mi-faser^14^, SUPER-FOCUS^15^, and eggNOG-Mapper^16^ rely on translated search against comprehensive protein databases, for example using DIAMOND^17^. Although these tools can be executed in combination, doing so yields independent taxonomic and functional profiles without per-sequence co-annotation.

HUMAnN integrated with MetaPhlAn represents the most widely adopted strategy for obtaining both community composition and functional potential from a single dataset^18^. It employs a tiered approach, in which MetaPhlAn first profiles species abundance using clade-specific marker genes, and the remaining unmapped reads are subjected to HUMAnN for functional annotation. This workflow is conceptually elegant and has proven effective across numerous studies. However, this architecture faces three practical constraints when applied to large-scale or complex environmental meta’omic data. First, MetaPhlAn offers high specificity and computational efficiency by targeting clade-specific genes, yet its sensitivity is inherently tied to the coverage of the marker gene database; in complex environmental samples, this can limit detection of taxa lacking representative markers^5,19^. Second, HUMAnN computes functional profiles against the UniRef database, which provides fine-grained resolution, but conversion to the widely used KEGG Orthology (KO) system introduces an additional mapping layer in which many UniRef entries lack one-to-one KO correspondences, so the converted profiles capture only part of the annotated signal. Third, the reliance on DIAMOND for functional annotation can become computationally intensive, potentially limiting scalability to very large datasets. Frameworks that instead achieve per-sequence co-annotation directly, such as MEGAN^20^, face this cost in its full form, as every read must be aligned against comprehensive protein databases.

Beyond these limitations, a more fundamental barrier to coupled taxonomic-functional profiling is the structural difference between taxonomic and functional databases^21^. Taxonomic classification benefits from the Last Common Ancestor (LCA) mechanism, which provides robust assignment for ambiguous sequences lacking sufficient phylogenetic signal or shared among closely related clades^22^. Functional profiling lacks an equivalent mechanism because KEGG and Gene Ontology (GO) employ network-like^23^ and directed acyclic graph structures^24^, respectively, precluding the hierarchical tree structures required for LCA. Consequently, *k*-mer-based taxonomic classifiers such as Kraken2, whose LCA assignment presumes a tree-structured reference, cannot be directly applied to functional annotation^21^. Existing functional classifiers such as Carnelian^25^ classify functions via compositional gapped *k*-mer features, and DIAMOND-based tools rely on best-hit assignment, both of which lack LCA-like mechanisms to resolve ambiguity among multiple equally good matches, reducing annotation confidence for sequences with shared domains or conserved regions^12,26^.

Meta’omic analysis can be conducted at three levels: reads, contigs, and MAGs^4^, yet existing tools typically operate on a single level (Figure 1a). Read-level profiling achieves maximal community coverage but suffers from reduced classification confidence due to short sequence length and limited phylogenetic signal, whereas contig-and MAG-level analyses offer higher annotation accuracy at the cost of incomplete community representation due to assembly^27^ and binning^28^ losses (Figure 1b and 1c). While multi-level integration has been shown to improve taxonomic profiling accuracy by balancing this inherent tradeoff^29^, systematic frameworks that jointly integrate taxonomic and functional information across reads, contigs, and MAGs remain undeveloped. In addition, cross-level comparison is impeded by inconsistent database usage. The Genome Taxonomy Database (GTDB)^30,31^ and the International Committee on Taxonomy of Viruses (ICTV)^32^ have recently undertaken systematic revisions of prokaryotic and viral classification systems, yet most existing tools lack native support for these updated standards or the capacity to simultaneously process both microbial and viral components. More critically, analytical level-specific database dependencies create classification incongruities, as GTDB is predominantly employed for MAG-level annotation (e.g., via GTDB-Tk^33^), whereas NCBI Taxonomy underpins contig-and read-level classification, yielding contradictory taxonomic assignments for identical taxa across analytical levels. This database fragmentation fundamentally undermines cross-level comparability and precludes integrative multi-level analysis.

**Figure 1.**
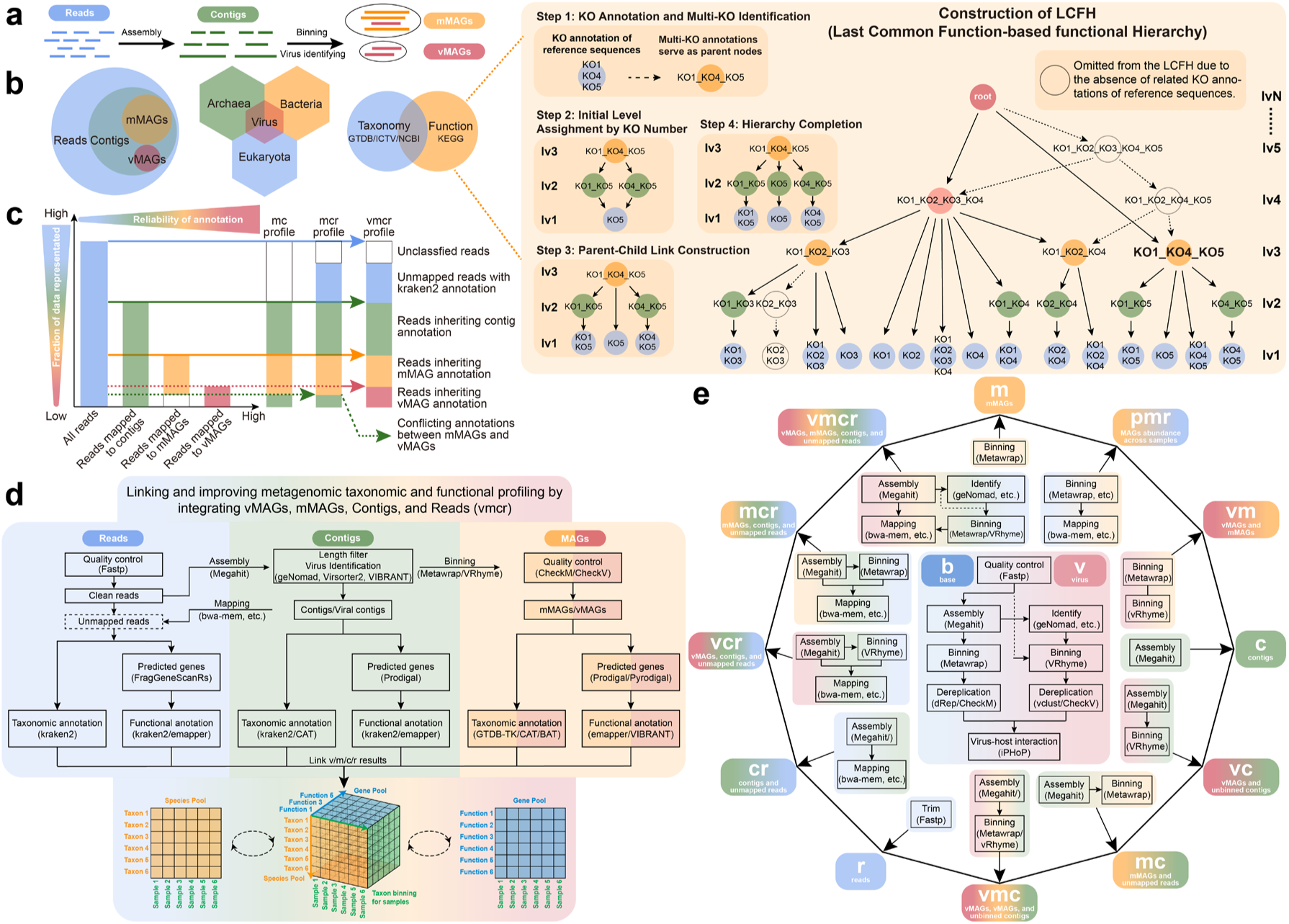
Overview of the Metalink workflow. **a**, Conventional assembly-and binning-based metagenomic analysis pipeline for microbial and viral communities. **b**, Metalink extends and improves conventional metagenomic analysis through three aspects: integration of reads, contigs, and metagenome-assembled genomes (MAGs); support for both the Genome Taxonomy Database (GTDB) and the International Committee on Taxonomy of Viruses (ICTV) databases; and rapid, accurate functional annotation via a custom-built functional database according to the Last Common Function-based functional Hierarchy (LCFH) framework (dashed box). **c**, Representative modes of Metalink illustrating the expansion of standard metagenomic analysis. Left, schematic bar plot showing the fraction of metagenomic data annotated as reads, contigs, microbial MAGs (mMAGs), and viral MAGs (vMAGs). Right, schematic overview of three analytical modes: Metalink-mc refines contig annotation using mMAG-derived results; Metalink-mcr further extends annotation by rapidly profiling unmapped reads; Metalink-vmcr integrates viral sequence identification to optimize Metalink-mc results, followed by read-level expansion analysis. **d**, Detailed workflow of Metalink-vmcr mode. **e**, Schematic summary of different Metalink modes. The base mode (Metalink-b) enables assembled contigs and binned mMAGs; Metalink-v enables viral prediction; and 12 additional modes, including Metalink-vmcr, support integrated taxonomic and functional analyses.

To enable direct linkage of species taxonomy to metabolic function at multiple levels, we developed Metalink with three main components: (i) The Last Common Function-based functional Hierarchy (LCFH) reorganizes multi-KO annotations into a tree-structured functional classification, enabling multiple taxonomic profilers like Kraken2 to perform reliable functional assignment with LCA-like resolution; (ii) A unified taxonomic framework integrates GTDB, ICTV, and NCBI for consistent annotation across MAG, contig, and read levels; (iii) Multi-level integration combines these three sequence types with overlap-based deduplication to balance accuracy and completeness. For functional profiling, Metalink exceeds both HUMAnN3 and eggNOG-Mapper in read-level assignment accuracy while operating approximately 30-fold and 1,000-fold faster, respectively, and its contig-level profiles remain highly consistent with eggNOG-Mapper. For taxonomic profiling, benchmarking on CAMI II datasets^4^ shows Metalink attains F1 score 0.955 and completeness 0.971, outperforming state-of-the-art taxonomic profilers. The joint output of functional and taxonomic profiles at multiple levels provides direct means to link species with their encoded functions.

## Results

### Overview of the Metalink workflow

Metalink implements an integrated strategy that functionally and taxonomically profiles metagenomic sequencing at MAG, contig, and read levels (Figure 1b and Fig. S1). Metalink provides twelve processing modes organized by sequence type inclusion and viral analysis scope. Except for read-only modes, all modes depend on the Metalink-base mode for contig assembly and mMAG binning, and all virus-related modes additionally depend on the Metalink-virus mode for viral prediction and viral binning workflows. For example, the vmcr (vMAGs-mMAGs-Contigs-Reads) mode implements the complete pipeline: clean reads are assembled into contigs, and contigs exceeding length thresholds are binned into microbial MAGs (mMAGs) and viral MAGs (vMAGs) (Figure 1d). Reads are mapped to contigs for abundance quantification, and unassembled reads are retained at the read level. Overlap-based deduplication resolves redundant annotations, with contigs incorporated into MAGs inheriting the MAG’s taxonomic and functional identities, and prokaryotic-viral MAG overlaps resolved with viral priority. This ensures each sequence contributes only once, yielding non-redundant community profiles. Taxonomic annotation employs Kraken2^8^ for reads, Kraken2 or CAT_pack^26^ for contigs, and GTDB-Tk^33^ or BAT^26^ for MAGs. Functional annotation employs eggNOG-Mapper or Kraken2 (translated search against the custom LCFH protein database) for reads, contigs and MAGs, and VIBRANT^34^ for viral sequences. Per-sequence taxonomic and functional information is aggregated to obtain per-organism gene family abundances in addition to community-level profiles. Other processing modes can be treated as simplified versions of the vmcr mode (Figure 1e). For example, the mcr (mMAGs-Contigs-Reads) mode excludes vMAG analysis, considering only mMAGs, contigs, and unassembled reads; the mc (mMAGs-Contigs) mode only uses mMAG information to optimize contig results; and the r (Reads) mode performs joint taxonomic and functional profiling using only read-level information.

### Taxonomic profiling performance of Metalink

To benchmark against existing tools, we evaluated Metalink’s performance using the NCBI taxonomic system snapshot consistent with CAMI II. We compared the comprehensive vmcr mode and the r mode against state-of-the-art tools (Figure 2). Benchmarking was performed using OPAL^35^ with metrics adopted from CAMI II. Pre-computed benchmark results for multiple tools were downloaded from the CAMI II website (https://cami-challenge.org/), and RAT^29^ was evaluated locally using identical methods. Metalink achieved the highest F1 score (vmcr mode, 0.955 ± 0.011) and completeness (0.971 ± 0.007), exceeding MetaPhlAn4, mOTUs, sylph^10^, and RAT at the genus level. The vmcr mode also exhibited high purity (0.939 ± 0.017), surpassing MetaPhlAn4, sylph, and RAT, though slightly below mOTUs. The r mode additionally showed the lowest L1 norm error (0.152 ± 0.030). Beyond the marine dataset, Metalink was tested on CAMI II plant-associated dataset with similarly superior performance (Extended Figure 1), indicating broad applicability across distinct environments.

**Figure 2.**
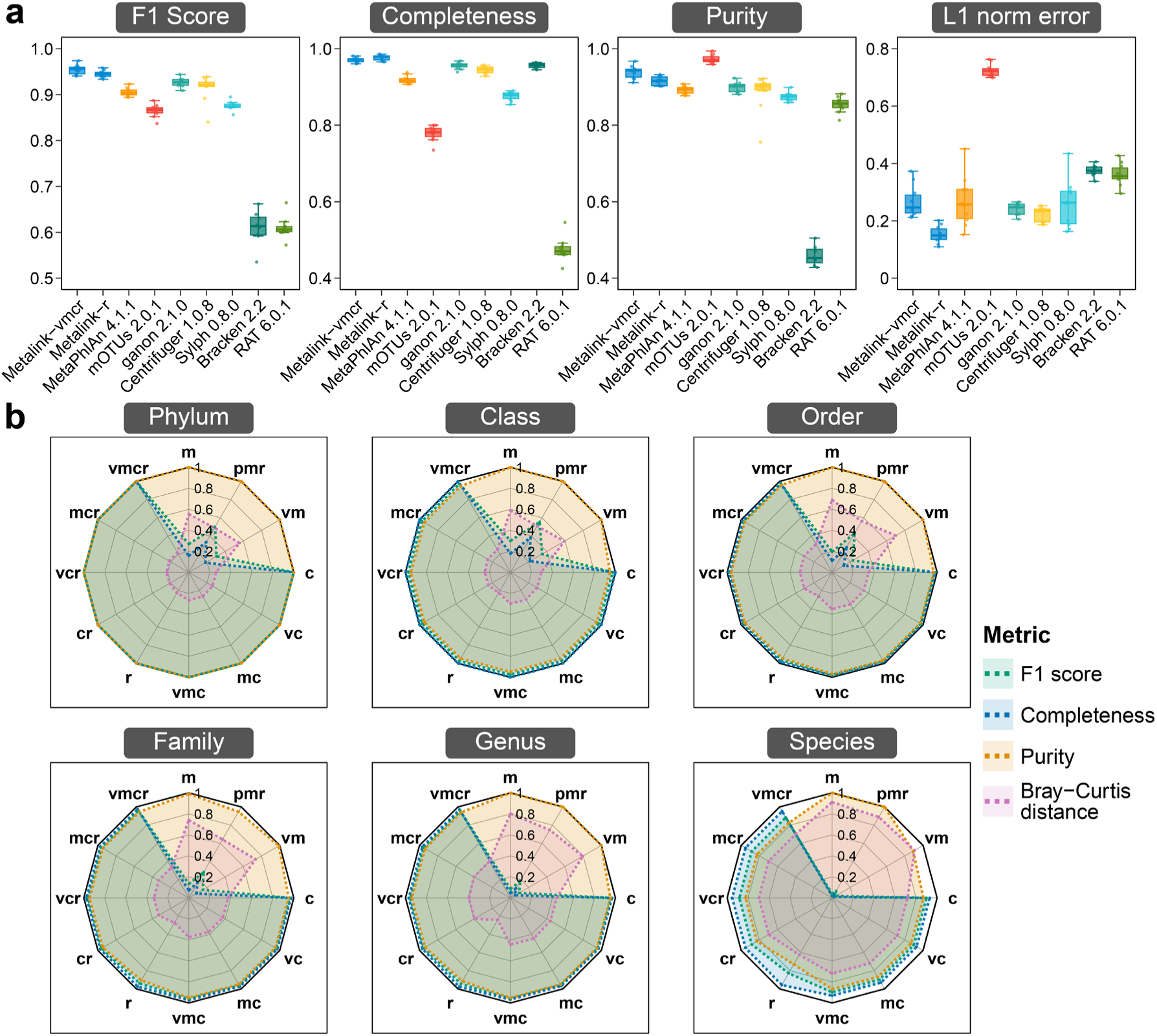
Taxonomic profiling performance of Metalink on CAMI II marine datasets. **a**, Benchmark comparison of Metalink-vmcr and Metalink-r modes against other tools at the genus level (n = 10 samples). **b**, Radar plots displaying taxonomic profiling results across various Metalink modes (n = 10 samples).

We further benchmarked different modes of Metalink across taxonomic levels. All modes showed better performance at higher levels, for example, order-level F1 score of vmcr was 0.973 ± 0.009, family level 0.960 ± 0.008, genus level 0.955 ± 0.011, and species level 0.885 ± 0.011. Modes including read-level analysis were compared, including vmcr and mcr, vcr (vMAGs-Contigs-Reads) and cr (Contigs-Reads), and r (Figure 2b). The vmcr and mcr modes achieved nearly identical performance, likely because CAMI II does not systematically assess viral components. The vcr and cr modes (lacking MAGs) were slightly weaker (0.954 ± 0.010) than vmcr and mcr at the genus level (0.955 ± 0.011). The r mode achieved the highest completeness (0.977 ± 0.007) but lowest purity (0.915 ± 0.012). Modes excluding read-level analysis were further compared, including vmc (vMAGs-mMAGs-Contigs), mc (mMAGs-Contigs), vc (vMAGs-Contigs), c (Contigs), vm (vMAGs-mMAGs), and m (mMAGs). The vmc and mc modes achieved F1 score 0.960 ± 0.009, slightly exceeding vc and c (0.959 ± 0.009). The vm and m modes showed the lowest completeness (0.050 ± 0.013) and highest purity (1.000 ± 0.000), opposite to the r mode. Finally, the pmr (Project-level pooled MAGs with Reads) mode outperformed the m mode. The pmr mode leverages a non-redundant MAG dataset (in this test, constructed from ten CAMI II marine samples), thereby increasing reference genome coverage. Its performance depends on the number and quality of MAGs, and scalability to larger sample collections remains to be evaluated.

As contig-level analysis is central to eight Metalink modes, we evaluated the impact of contig classification method and length threshold on Metalink taxonomic profiling performance (Extended Figure 2). Kraken2 outperformed CAT_pack across all lengths and modes, but the magnitude of improvement depended on contig length and mode configuration. At 1000 bp, Kraken2 moderately exceeded CAT_pack (vmcr F1 score 0.954 ± 0.009 versus 0.795 ± 0.021), but the gap widened substantially at 300 bp (0.955 ± 0.011 versus 0.469 ± 0.020). This likely reflects CAT_pack’s reliance on multi-open reading frame (ORF) voting^26^, which is severely weakened when 300 bp contigs contain few ORFs, whereas Kraken2’s *k*-mer voting remains effective on short contigs. The inclusion of read-level analysis differentially affected the two methods. In read-inclusive modes (vmcr, mcr, vcr, cr), Kraken2 performed similarly at 300 bp and 1000 bp (vmcr L1 norm error 0.372 ± 0.049 versus 0.373 ± 0.051), whereas CAT_pack was substantially better at 1000 bp (vmcr L1 norm error 0.462 ± 0.069 versus 0.648 ± 0.038). In read-exclusive modes (vmc, mc, vc, c), Kraken2 remained superior at 300 bp (vmc L1 norm error 0.444 ± 0.053 versus 0.689 ± 0.050), while CAT_pack performed poorly at both lengths.

### Taxonomic binning performance of Metalink-r

We benchmarked Metalink’s taxonomic binning performance for GTDB (prokaryotes) and ICTV (viruses) components of the integrated taxonomic framework (Fig. S1), as these represent the primary advances over conventional NCBI-based pipelines^5,36^. Based on version splits of GTDB and ICTV databases, genomes present in both older and newer versions were designated as the known genomes set, while genomes newly added in the newer version were designated as the unknown genomes set (Extended Figure 3). Known genomes were used to construct the Metalink taxonomic database. Both known and unknown genomes were randomly fragmented to simulate metagenomic reads for HiSeq, NextSeq, and NovaSeq platforms. Performance of both Metalink and MetaPhlAn4 was evaluated on the known and unknown sets (Extended Figure 4), with the benchmark of Metalink reflecting Kraken2-based detection of reads across entire genome regions, whereas that of MetaPhlAn4 reflects detection accuracy for reads mapped to marker genes.

For known prokaryotes, Metalink outperformed MetaPhlAn4, with genus-level F1 scores of 0.9510-0.9642 versus 0.8755-0.8805. Purity at the genus level was also higher (0.9668-0.9758 versus 0.8923-0.8962). For unknown prokaryotes, Metalink performance decreased substantially, with genus-level F1 scores of 0.3573-0.6000 versus MetaPhlAn4’s 0.8975-0.9178. Several factors may account for this difference. First, although MetaPhlAn4 provides multiple database releases^10^, its marker set is organized around SGBs that do not correspond one-to-one to versioned GTDB genomes, so the extent to which the ‘unknown’ genomes in our benchmark are already represented in its marker set cannot be assessed exactly. In contrast, Metalink employs version-traceable database construction (GTDB R220 and ICTV MSL38 genomes for construction, versus genomes unique to R226 and MSL40 for unknown-species benchmarking), enabling strict control over reference genome composition. Second, alignment-based profiling (MetaPhlAn4) may exhibit greater sensitivity than *k*-mer-based classification for highly divergent unknown sequences under the parameter settings evaluated here, which may partly favor MetaPhlAn4 in the unknown-genome comparison. Additionally, Metalink extends taxonomic identification to viruses, which MetaPhlAn4 currently does not.

### Construction of hierarchical functional classification system

Direct construction of a tree-structured classification system from arbitrary gene family permutations is computationally infeasible. We therefore treated all protein annotations from genomes in the integrated taxonomy database as a representation of the known protein space, from which we captured observed KO annotation combinations. Accordingly, gene prediction yielded 424,152,473 protein sequences, from which eggNOG-Mapper functional annotation produced 45,231 non-redundant KO annotation combinations comprising 16,710 single-KO and 28,521 multi-KO annotations. These multi-KO annotations represent similar KOs co-assigned to individual sequences, indicating functional redundancy or shared conserved domains among these KO clusters.

By utilizing this annotation information, we designed LCFH (Figure 1b), a tree-structured functional classification where each node represents a set of KOs and parent-child relationships are established by strict set inclusion with the unique parent rule (a node is assigned to a higher-level parent only when exactly one containing node exists; otherwise it retains its identity) (see Methods). This resolves the combinatorial explosion of arbitrary node permutations while enabling *k*-mer-based search tools to perform functional annotation with LCA-like resolution^21^. Statistical analysis of all unique KO annotations revealed that over 99% contain seven or fewer KOs (Extended Figure 5). LCFH was therefore set to seven levels, maintaining simplicity while matching the number of taxonomic levels in the GTDB.

Because different LCFH levels lack explicit similarity thresholds, potentially causing uncertainty in practical application, we designed the Similarity Clustering with Frequency-filtered functional Hierarchy (SCFH) as an approximate positioning system (Extended Figure 6). In SCFH, all annotated protein-coding sequences were clustered by MMseqs2^37^ at given similarity thresholds, yielding seven levels with representative sequences retained at each level: KO_100% (201,416,117 sequences), KO_90% (195,955,152), KO_70% (45,875,212), KO_50% (10,925,368), KO_40% (3,746,910), KO_30% (1,170,864), and KO_20% (534,810). Representative KO sets were then selected from each cluster at cumulative frequency thresholds of 2%, 5%, 10%, 20%, 30%, 40%, and 50% (the resulting hierarchies hereafter denoted SCFH_2 through SCFH_50) to define the most representative node names at each level (Fig. S2-S5).

### Benchmark of hierarchical functional classification system

We benchmarked LCFH and SCFH using the split-database approach, with SCFH also serving as an anchor to approximately position LCFH. Old GTDB genomes provided the known gene set for Metalink functional database construction and benchmarking, while newer genomes provided the unknown gene set for benchmarking (Figure 3a). All genes from both sets were fragmented using HiSeq, NextSeq, and NovaSeq error models to generate test reads (Extended Figure 3). Per-read annotation accuracy was assessed at each functional classification level, using the eggNOG-Mapper annotation (eggNOG v5.0.2^16^) of the source gene prior to fragmentation as the reference.

**Figure 3.**
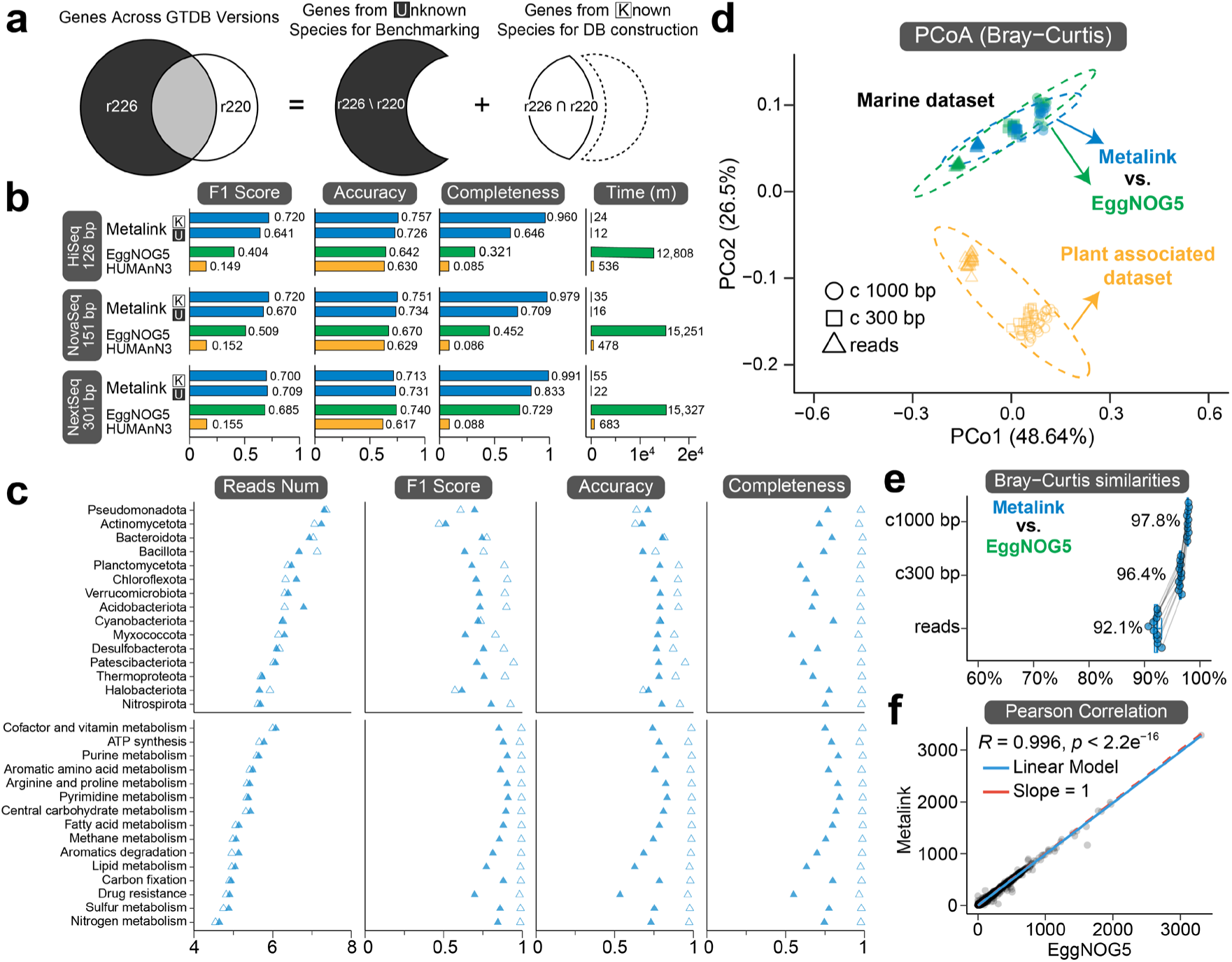
Functional binning performance of Metalink. **a**, Benchmarking of functional binning of Metalink-r using a split-database approach to distinguish functions from known and unknown species. **b**, Functional binning performance of Metalink-r compared to eggNOG-Mapper (EggNOG5) and HUMAnN3 across different sequencing platforms. **c**, Functional binning performance of Metalink-r across taxonomic groups and metabolic pathways using simulated NovaSeq reads. **d**, Principal coordinate analysis (PCoA) based on Bray-Curtis distances comparing functional binning consistency between Metalink and eggNOG-Mapper on the CAMI II marine dataset (n = 10 samples). Three data types were evaluated: clean reads, contigs retained at 300 bp and 1,000 bp length thresholds. The CAMI II plant-associated dataset served as a control group (n = 21 samples). **e**, Bray–Curtis similarity between Metalink and eggNOG-Mapper functional binning results for the CAMI II marine dataset at three sequence types: raw reads, 300 bp contigs, and 1,000 bp contigs. f, Consistency between Metalink and eggNOG-Mapper annotations for a marine metagenomic sample from CAMI II at the contig 1,000 bp level. Each point represents a KO, with the x-and y-axes showing the mean coverage of that KO annotated by different software, respectively.

We evaluated F1 score, classified percent, and node name complexity for each functional classification level (Extended Figure 7). In LCFH and SCFH, average KO count per node increased from lower to higher levels (lv1 to lv7), with F1 score and classified percent showing a clear negative correlation. For example, LCFH lv1 and SCFH KO_100% achieved the highest F1 score and lowest classified percent within their respective hierarchies.

We then used the product of F1 score and classified percent as an integrated benchmark metric (Extended Figure 8). Level 1 was similar across classification systems, largely because all methods employed identical lv1 (or KO_100%) KO annotations and reference sequences. The highest integrated score was achieved by the default LCFH lv1 level (28.81). Higher levels showed greater divergence between LCFH and SCFH. The performance decline from LCFH lower to higher levels (lv1/lv7) was substantially smaller than that of SCFH, though this may partly reflect non-correspondence between level definitions. In fact, LCFH lv1 to lv7 approximately matched SCFH KO_100% to KO_50% in both F1 score and classified percent. For example, LCFH lv7 and SCFH_40 at the KO_50% level showed comparable F1 scores.

### Parameter sweeps for Metalink functional database

We systematically scanned the Kraken2 database construction parameters—*k*-mer length (*k*), minimizer length (*ℓ*), and minimizer spacing (*s*)—for nucleotide (ffn) and protein (faa) databases against LCFH to identify defaults optimizing the trade-off between accuracy, completeness, and computational cost. Testing on unknown species across all three error models showed that protein databases substantially outperformed nucleotide databases in F1 score, sensitivity, and completeness. For example, top-performing protein databases on the NovaSeq platform achieved F1 scores of 0.656-0.670 versus 0.511-0.523 for nucleotide databases (Extended Figure 9). These results indicate that protein-based Metalink databases generalize substantially better than nucleotide databases to novel species. Testing on known species showed protein databases were slightly weaker than nucleotide (F1 score 0.689-0.720 versus 0.748-0.753), but with substantially shorter runtime (27-35 min versus 84-144 min) and lower peak memory (63.7-66.0 GB versus 168.2-276.9 GB). The performance pattern of nucleotide and protein databases reflects a fundamental trade-off between phylogenetic resolution and functional generalizability, as nucleotide *k*-mers capture codon usage bias and synonymous substitution patterns that are highly characteristic of closely related, known lineages within the reference database, whereas protein *k*-mers retain discriminative signal across broader taxonomic distances where nucleotide sequences have diverged beyond reliable matching. Considering generalizability requirements for samples dominated by potentially novel species, we selected the parameter combination (*ℓ* = 17, *k* = 14, and *s* = 1) that maximized F1 score on unknown genomes, which also achieved optimal or near-optimal performance across accuracy, purity, specificity, and completeness.

### Functional binning performance of Metalink-r

Using the LCFH and the parameter combination (*ℓ* = 17, *k* = 14, and *s* = 1), we compared Metalink-r against eggNOG-Mapper and HUMAnN3 for per-read functional assignment accuracy (Figure 3b, and Extended Figure 3). For KO annotations of simulated reads across error models, Metalink achieved F1 scores of 0.641-0.720, exceeding eggNOG-Mapper (0.404-0.685) and HUMAnN3 (0.149-0.155). Completeness was also substantially higher (0.646-0.991 versus 0.321-0.729 and 0.085-0.088). The lower F1 scores and completeness of HUMAnN3 in this evaluation largely reflect the indirect UniRef-to-KO conversion: a substantial proportion of UniRef entries lack one-to-one KO correspondences, so the converted KO profiles capture only part of the annotated signal. We note that HUMAnN3 is designed for community-level pathway quantification rather than per-read KO assignment. Notably, eggNOG-Mapper accuracy and completeness approached Metalink as read length increased, indicating Metalink’s superior performance on shorter sequences relative to eggNOG-Mapper. In addition, Metalink runs approximately 30-fold faster than HUMAnN3 and three orders of magnitude faster than eggNOG-Mapper, with higher peak memory (62.5–75.6 GB) than HUMAnN3 (30.1–35.8 GB) and eggNOG-Mapper (17.8–18.6 GB), though remaining practical on standard high-memory servers (Figure 3b and Extended Figure 9).

We also assessed per-phylum and per-pathway functional binning accuracy of Metalink. Per-phylum accuracy was generally high across the top 15 representative phyla, comprising 5.6 × 10^5^ to 2.3 ×10^7^ sequences and approximately 90% cumulative abundance of all tested sequences (Figure 3c), though *Actinomycetota* exhibited notably lower F1 score, suggesting phylum-specific characteristics or higher within-phylum functional divergence, and *Myxococcota* showed markedly lower completeness for unknown species, indicating substantial uncharacterized functions in this predatory group not captured by current databases^38^. Per-pathway accuracy was similarly high for 15 representative pathways, comprising 3.5 × 10^4^ to 1.2 × 10^6^ sequences and approximately 4% cumulative abundance. with drug resistance as an exception, showing lower F1 score and completeness for unknown species than for known species, reflecting substantial divergence of resistance genes between characterized and uncharacterized species.

### Functional profiling performance of Metalink

We evaluated the concordance between read-level and contig-level functional profiles generated by Metalink, using eggNOG-Mapper as the reference standard (Figure 3d and 3e). PCoA of Bray-Curtis distances clearly separated CAMI II marine and plant-associated datasets regardless of sequence type or software method (Figure 3d). For the marine dataset, contig 300 bp functional profiles are thought to represent a balance between accuracy and completeness, mirroring the taxonomic profiling pattern where contig 300 bp taxonomic profiles were closer to the reference than contig 1000 bp (Extended Figure 2). Metalink read-level profiles clustered closer to contig 300 bp than eggNOG-Mapper results, indicating closer agreement of Metalink’s read-level profiles with the contig-level reference, consistent with the functional binning benchmark.

We then compared Metalink and eggNOG-Mapper across sequence types for CAMI II marine datasets (Figure 3e). Bray-Curtis similarity between the two tools was highest at contig 1000 bp (97.8%), slightly lower at contig 300 bp (96.4%), and lowest at read level (92.1%), consistent with the expectation that longer sequences provide stronger signal for eggNOG-Mapper. Per-KO abundance comparison showed high consistency between Metalink and eggNOG-Mapper at contig 1000 bp (Figure 3f and Fig. S6). Overall, Metalink provides higher accuracy and substantially faster speed at read level, while maintaining high consistency with eggNOG-Mapper at contig level.

### Integrated taxonomic and functional profiling of metagenomic and metatranscriptomic datasets using Metalink

To demonstrate the joint analysis capability for taxonomic and functional profiling of metagenomic and metatranscriptomic samples, we applied Metalink to two datasets: (1) a metagenomic dataset comprising 10 samples from each of 29 distinct environments (Supplementary Data 1), and (2) a metatranscriptomic dataset of 108 seawater samples from the TARA Oceans expedition^39,40^ (Supplementary Data 2).

Read-level functional profiling showed that microbial communities across diverse environments, including surface seawaters, deep-sea hydrothermal vents, cold seeps, terrestrial hot springs, soil, gut, and food systems, exhibited conserved functional compositions, with core metabolic pathways such as amino acid metabolism, carbohydrate metabolism, and energy metabolism being dominant (Fig. 4a). In contrast, taxonomic profiles varied across environments. Although *Pseudomonadota* was the predominant phylum in most environments, its relative abundance varied considerably. Even within different microhabitats of deep-sea hydrothermal vents, taxonomic profiles varied substantially. For instance, plumes and ambient seawater both harbored taxa such as SAR324, whereas subsurface fluids enriched in *Bacteroidota* showed greater compositional similarity to chimney communities.

**Figure 4.**
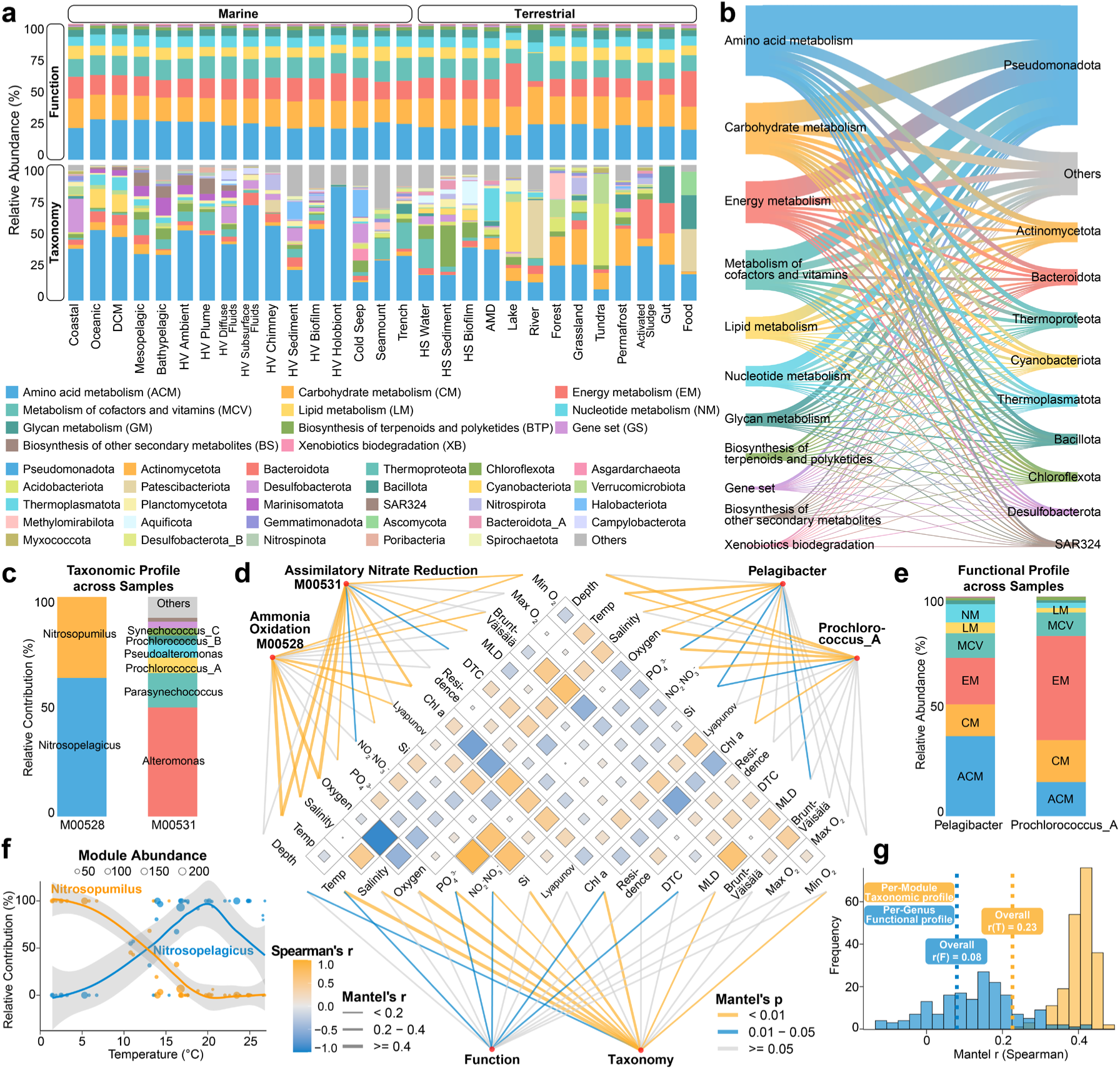
Functional and taxonomic profiles, and their coupling, across marine and terrestrial metagenomics and metatranscriptomes. a, Mean relative abundance of KEGG metabolic pathways and the top 29 most abundant phyla across marine and terrestrial environments. Each bar represents the average abundance of 10 samples per environment. Colors indicate metabolic categories and phyla as defined in the legend. **b**, Sankey diagram linking metabolic pathways to phyla based on aggregated mean abundance across the 29 environments. Line width reflects the magnitude of taxonomic contribution to each pathway. **c**, Taxonomic composition of two selected metabolic modules (ammonia oxidation, M00528; assimilatory nitrate reduction, M00531) based on mean relative abundance across 108 TARA Ocean metatranscriptomic samples. **d**, Mantel test correlations between transcripts and environmental factors across 95 of 108 TARA Ocean metatranscriptomic samples with available environmental metadata at three levels: per-module taxonomic profiles (upper left), per-genus functional profiles (upper right), and overall taxonomic and functional profiles (lower). The heatmap shows pairwise Spearman correlations among environmental factors. Line width and color indicate the Mantel *r* statistic and significance, respectively. **e**, Functional composition of two selected genera (*Pelagibacter* and *Prochlorococcus_A*) based on mean relative abundance across TARA Ocean samples. **f**, Relative contribution of *Nitrosopumilus* and *Nitrosopelagicus* to ammonia oxidation (M00528) across a temperature gradient. Points represent individual samples, with point size proportional to the total abundance of M00528. Solid lines indicate LOESS-smoothed trends with 95% confidence intervals (shaded areas). **g**, Distribution of Mantel *r* values from per-module taxonomic profiles and per-genus functional profiles correlated with environmental factors across TARA Ocean samples. Dashed lines indicate the Mantel r values for the overall genus profile and module profile across all samples. DCM, deep chlorophyll maximum. HV, hydrothermal vent. HS, hot spring. AMD, Acid Mine Drainage.

Furthermore, a key advantage of Metalink lies in its support for establishing quantitative linkages between taxa and metabolic functions. At the phylum level, we observed extensive decoupling between functional and taxonomic profiles. Both core metabolic pathways (e.g., amino acid metabolism, carbohydrate metabolism, and energy metabolism) and specialized functional pathways (e.g., biosynthesis of terpenoids and polyketides, and xenobiotics biodegradation) were contributed by multiple phyla (Fig. 4b). This broad taxonomic participation indicates that metabolic functions are distributed across diverse microbial lineages rather than confined to specific taxa, though the degree of functional redundancy varies across pathways. For example, in surface seawaters, ammonia oxidation (M00528) was dominated by two genera, *Nitrosopumilus* and *Nitrosopelagicus*, compared with the multiple contributors observed for assimilatory nitrate reduction (M00531) (Fig. 4c). Additionally, the cross-environment stability of functional profiles does not imply functional consistency across taxa. For example, *Prochlorococcus_A* exhibited substantially higher functional expression related to energy metabolism than *Pelagibacter* (Fig. 4e).

We next used Metalink’s joint outputs to examine microbiome–environment associations at two levels: the taxonomic composition of functionally contributing taxa, and per-genus functional profiles (Fig. 4d-g). As highlighted in previous studies, aggregated taxonomic or functional profiles at the community level are tightly associated with environmental factors based on Mantel tests^41^. We selected two nitrogen metabolism modules to demonstrate the linkage between the taxonomic composition of functionally contributing taxa and environmental factors (Fig. 4c and d). The species composition of M00528 (ammonia oxidation) was strongly correlated with environmental factors such as temperature and salinity, suggesting that these factors are key determinants of the contributing taxa (e.g., *Nitrosopumilus* versus *Nitrosopelagicus*). Temperature-correlated niche partitioning between these two ammonia-oxidizing archaea was further supported by their contrasting relative contributions to M00528 across a temperature gradient (Fig. 4f). Meanwhile, contributing taxa of this module showed no significant correlation with nitrate/nitrite concentrations, suggesting no detectable selective pressure from product accumulation at the resolution of our data. In contrast, variations in nitrate/nitrite were significantly correlated with shifts in the taxa contributing to M00531 (assimilatory nitrate reduction), implicating substrate concentration as a key driver of taxonomic turnover. We further selected the two most abundant marine genera, *Prochlorococcus_A* and *Pelagibacter*, to illustrate differential functional responses to environmental factors. The results showed that nitrate/nitrite and phosphate were more strongly associated with functional profile variation in *Prochlorococcus_A* than in *Pelagibacter*, suggesting that nutrient limitation may be a primary driver of functional expression in *Prochlorococcus_A* (Fig. 4d and e). In contrast, *Pelagibacter* functional expression showed weaker associations with nitrate/nitrite and phosphate, suggesting that this heterotroph is less responsive to inorganic nutrient fluctuations.

In summary, Metalink provides an extensible framework for integrating taxonomic and functional profiles, enabling direct tests of how microbiome composition and function covary with environmental contexts. It should be cautioned that the specific associations reported in these case studies are statistical in nature, and their causal relationships require further validation through rigorous experimental approaches such as cultivation experiments or genetic manipulation.

## Discussion

The Last Common Ancestor mechanism in taxonomic profiling provides robust assignment for ambiguous sequences that lack sufficient phylogenetic signal or are shared among closely related clades^22^. Functional profiling has lacked this equivalent. LCFH fills this gap by capturing KO co-assignment patterns from 424 million protein sequences, using multi-KO annotations as internal nodes to construct a data-driven tree-like hierarchy. Unlike ontology-driven frameworks, LCFH derives hierarchical relationships from empirical co-annotation patterns, which reflect evolutionary conservation of protein domain architectures. This structure enables *k*-mer-based search tools— originally designed for taxonomic classification—to perform functional annotation with LCA-like resolution, improving reliability for ambiguous sequences. SCFH complements this framework as an approximate positioning system, defining quantifiable functional levels based on sequence similarity thresholds to provide empirical anchoring for LCFH. LCFH requires periodic updating, analogous to GTDB, which releases new versions periodically to expand coverage. The current LCFH is constructed based on KO annotations. Future iterations may integrate broader annotation sources such as GO or Enzyme Commission classification systems^21^.

The standardization of taxonomic frameworks has become increasingly important, yet the contradictions arising from different databases used at different analytical levels represent not merely technical inconveniences but conceptual barriers. GTDB is predominantly used at the MAG level, while NCBI is widely employed at the contig and read levels. This makes it difficult to systematically verify whether classification results for the same dataset are consistent across levels. Metalink’s integrated taxonomic system guarantees that a taxonomic unit carries the same identifier and lineage whether it is identified from a read, contig, or MAG. This makes cross-level consistency a measurable analytical property. In addition, the integration of the ICTV viral taxonomic system is particularly critical, enabling viruses and prokaryotes to be processed within the same pipeline without requiring separate analytical workflows.

Building upon this foundation of taxonomic and functional profiling, multi-level integration of Metalink treats MAGs, contigs, and reads as parallel information sources, consolidating them into a unified taxonomic-functional profile through overlap-based deduplication. Existing methods typically operate at a single analytical level^9,14,25^. Read-level methods achieve maximal coverage but carry limited information per sequence, while assembly^27^-and binning^28^-based methods offer higher accuracy but discard unassembled or unbinned reads. Metalink instead propagates MAG-level assignments to their constituent contigs (for both mMAGs and vMAGs) and retains unassembled reads, so that the accuracy gains at higher levels are not paid for with coverage loss. While benchmarking on CAMI II datasets showed comparable performance across modes due to the high assembly efficiency of these simulated communities, multi-level integration of Metalink is expected to provide greater benefits for complex environmental samples with lower assembly rates, where read-level profiling becomes essential to capture community members discarded during assembly or binning. Critically, Metalink not only optimizes accuracy and completeness of taxonomic classification but also incorporates functional classification into the same framework at substantially reduced computational cost compared to the taxonomic profiler RAT, enabling joint multi-level analysis of function and taxonomy (Extended Figure 10).

Metalink’s multi-level joint output, where the same set of sequences co-annotated with both taxonomic and functional annotations, enables direct quantification of which microorganisms contribute to which metabolic functions, thus addressing a fundamental limitation of overall taxonomic and functional profiles. For example, overall taxonomic profiles can indicate that *Nitrosopumilus* abundance correlates with temperature, but reveal nothing about which metabolic functions it contributes, because the marker genes used for species identification are not functionally annotated. Conversely, overall functional profiles can indicate that ammonia oxidation gene abundance correlates with temperature, but cannot identify the executing microorganisms, because functional assignments are made independently of taxonomic provenance. Metalink assigns both profiles and provides detailed data on structural variations of functionally contributing assemblages across samples, or differential expression of gene complements within specific taxa. This advances analysis from community-level associations to species-resolved links between metabolic functions and environmental conditions.

Several limitations remain in the current Metalink implementation. The reference-dependency of LCFH means that genes without KO annotation or KO combinations not observed in reference genomes cannot be classified. Viral contigs may also be incompletely recovered by the adopted prediction tools, leading to an underestimation of viral MAG recovery. The eukaryotic component currently comprises only hundreds of genomes from NCBI RefSeq, a substantial gap compared to the hundreds of thousands of prokaryotic genomes in GTDB; accommodating larger and more numerous eukaryotic genomes may therefore require the integration of more efficient tools. In addition, to maintain a consistent taxonomic backbone across GTDB, ICTV, and NCBI, the unified framework retains only the seven principal ranks shared by these systems, so intermediate groupings used by individual sources are collapsed and cannot be resolved in downstream analyses. Finally, Metalink’s databases entail higher peak memory than some comparators (62.5– 75.6 GB in read-level functional binning), though this remains practical on standard servers, and the scalability of the pmr mode to much larger MAG collections remains to be evaluated.

In summary, Metalink restructures metagenomic taxonomic and functional analysis from independent tasks into joint outputs through three components, the LCFH functional hierarchy, an integrated taxonomic framework, and multi-level integration. Metalink improves overall performance in both taxonomic and functional profiling. More critically, it shifts the analytical focus from cataloging what functions are present in the community to determining which species perform which functions under which environmental conditions, making species–function associations directly observable.

## Methods

### Metalink pipeline

Metalink implements multiple processing modes that integrate MAG, contig, and read-level analyses through a Snakemake-based workflow with robust dependency tracking, dry-run preview, and resume capability. Here we describe the vmcr mode; other modes (vcr, mcr, cr, vmc, mc, vc, c, r) are simplified variants thereof. The workflow supports flexible entry points via the --skip-level parameter, allowing users to inject pre-trimmed reads (--skip-level 1), pre-assembled contigs (-- skip-level 2), or pre-binned MAGs (--skip-level 3) to bypass upstream steps. It also automatically adapts to diverse sequencing platforms through the --read-type parameter that selects appropriate trimming and assembly tools.

In the vmcr mode, raw reads are first trimmed using fastp (for short reads) or fastplong (for long reads). The choice of trimming and assembly software is determined automatically by the -- read-type parameter, which distinguishes paired-end, single-end, and long-read inputs (PacBio raw/corrected/HiFi; ONT raw/corrected/high-quality). For short-read metagenomes, reads are assembled using MEGAHIT v1.2.9^27^ with the meta-sensitive parameter or metaSPAdes v4.3.0^42^; for metatranscriptomes, rnaSPAdes^43^ is used; for long-read metagenomes, metaFlye v2.9.6^44^ is employed with parameters corresponding to the specific read type. Assembled contigs are binned using MetaWRAP v1.3.2^28^ (including CONCOCT v1.0.0^45^, MaxBin2 v2.2.6^46^, and MetaBAT2 v2.12.1^47^) with a minimum contig length threshold of 1,500 bp. Metalink implements a modified ViWrap v1.3.1^48^ framework for viral identification and quality control, with three configurable strategies: (i) geNomad v1.12.0^49^; (ii) intersection of VirSorter2 v2.2.4^50^ (--include-groups dsDNAphage,NCLDV,RNA,ssDNA,lavidaviridae) and VIBRANT v1.2.1^34^; or (iii) union of geNomad and the VirSorter2–VIBRANT intersection. Prophage information is predicted using ViWrap scripts. Lytic and lysogenic viruses are binned into vMAGs using vRhyme v1.1.0^51^. The quality of mMAGs and vMAGs is assessed using CheckM v1.2.5^52^, CheckM2 v1.1.0^53^, and CheckV v1.0.3^54^, respectively. Clean reads are mapped to contigs using minibwa v0.4^55^ (or alternatively BWA-MEM2 v2.2.1^56^, BWA mem v0.7.17^57^ and StrobeAlign v0.14.0^58^) and SAMtools v1.21^59^ for abundance quantification, and unassembled reads are retained at the read level. Unassembled reads are merged using fastp v0.23.4^60^ with the --merge option. All unassembled reads (merged paired-end reads and unmerged singletons) are subjected to ORF prediction using FragGeneScanRs v1.1.0^61^. Contigs incorporated into MAGs inherit the MAG’s taxonomic and functional identities; where prokaryotic and viral MAGs overlap, viral assignments take priority. Taxonomic annotation employs Kraken2 v2.1.3^8^ for reads, Kraken2 or CAT_pack v6.0.1^26^ for contigs, and GTDB-Tk v2.4.1^62^ or BAT from CAT_pack for MAGs. Functional annotation employs eggNOG-Mapper v2.1.12^16^ or Kraken2 (translated search) for read, contig, and MAG-level ORFs, and VIBRANT for viral ORFs. Per-sequence taxonomic and functional information is aggregated to obtain per-taxa gene family abundances in addition to community-level profiles. The custom R package mclink v1.3.1 was used to calculate metabolic pathway abundance and completeness.

Additional modes include m, v, vm, and pmr modes with specialized workflows. In the m and pmr modes, mMAGs are dereplicated using dRep v3.4.0^63^ with a similarity threshold (-sa) of 0.95. Prodigal v2.6.3^64^ with the parameter-p single is used to predict MAG ORFs. In the v mode, vMAGs are clustered using vclust^65^, and viral host prediction is performed using iPHoP v1.4.1^66^. The vm mode runs the complete v and m workflows and can further supply iPHoP with the dereplicated mMAGs produced by the workflow itself for host prediction. The pmr mode couples project-level MAG analysis with per-sample read quantification, with the MAG source selected via --mags m|v|vm (default m): mMAGs only (m), vMAGs only (v), or mMAGs and vMAGs merged into a single reference (vm). minibwa (or BWA-MEM2, BWA mem and StrobeAlign) and SAMtools are used to calculate mapping information of MAGs in each sample. CoverM v0.7.0^67^ in genome mode is used to summarize MAG abundance per sample, and in contig mode to summarize contig abundance per sample.

### Metalink taxonomic classification system construction

We constructed an integrated taxonomic classification system comprising three components: GTDB^30,31^ R226 for prokaryotic genomes, ICTV^32^ MSL40 for viral genomes, and NCBI Taxonomy (RefSeq Release 221) for microeukaryotic classification. Classification files including genome accession numbers and corresponding taxonomy were downloaded from GTDB r226 (https://data.ace.uq.edu.au/public/gtdb/data/releases/release226/226.0/), ICTV MSL40 (https://ictv.global/sites/default/files/MSL/ICTV_Master_Species_List_2024_MSL40.v1.xlsx), NCBI Taxonomy and Assembly databases (https://ftp.ncbi.nlm.nih.gov/pub/taxonomy/taxdump.tar.gz and https://ftp.ncbi.nlm.nih.gov/genomes/genbank/assembly_summary_genbank.txt). Taxonkit^68^ was used to generate stable identifiers from hierarchical names: taxid for taxonomic nodes (from taxon names and genome accessions) and funid for functional nodes (from KO annotations), ensuring version traceability and identifier consistency across database updates (e.g., names.dmp and nodes.dmp files).

Genome accession numbers were standardized to GCA prefixes without version suffixes to maintain stable taxid assignments while allowing genome updates across versions. For consistency, only the following ranks were retained: domain (*Bacteria*, *Archaea*, *Eukaryota*, and *Viruses*), phylum, class, order, family, genus, and species. To distinguish identical names at different taxonomic ranks, rank prefixes were added to taxon names during taxid generation, for example p UBA3054, c UBA3054, o UBA3054 for the placeholder’s name UBA3054 occurring at multiple ranks. Intermediate rank gaps can cause erroneous taxid assignments when identical abbreviated names appear under different higher-level contexts. For instance, c Caudoviricetes;o Autographivirales;f;g Pelagivirus and c Caudoviricetes;o;f;g Detrevirus would be incorrectly merged at the family rank if assigned solely by f. To prevent this, missing ranks were temporarily named using their parent rank information during database construction, for example f unclassified o Autographivirales family for the undefined family under o Autographivirales.

### Metalink taxonomic database construction

We constructed Metalink taxonomic databases using genomes from the integrated taxonomic classification system comprising GTDB, ICTV, and RefSeq microeukaryotes. Viral and plasmid regions were predicted and masked from prokaryotic and microeukaryotic genomes using geNomad to obtain clean prokaryotic and eukaryotic genome sequences. For Kraken2-based taxonomic databases, Kraken2-build was used with the established parameters (*ℓ* = 35, *k* = 31, and *s* = 7)^8^, which were likewise optimal for nucleotide databases in our functional-database parameter sweep (Extended Figure 9).

For CAT_pack-based taxonomic databases, all protein sequences were first dereplicated at 100% identity using MMseqs2. Mapping relationships between all protein sequences and dereplicated sequences were then used to assign taxid identifiers to non-redundant sequences, ensuring that the database uses non-redundant protein sequences while allowing these sequences to be mapped to different hosts sharing identical sequences. The command for constructing CAT_pack taxonomic databases was CAT_pack prepare.

### Taxonomic profiling benchmark

To enable comparable benchmarking against existing tools, we reconstructed a reference database using the NCBI taxonomy snapshot provided by CAMI II^4^ (https://openstack.cebitec.uni-bielefeld.de:8080/swift/v1/CAMI_2_DATABASES/). We used this database to benchmark twelve Metalink modes on the CAMI II metagenomic datasets. Additionally, we downloaded benchmark results for commonly used tools from the CAMI II website. As RAT lacks CAMI II benchmark results, we performed local testing using the same NCBI taxonomy snapshot as employed for Metalink. Performance was evaluated using the OPAL workflow^35^, generating metrics including F1 score, purity (precision), completeness (recall), L1 norm error, and Bray-Curtis distance.

### Taxonomic binning benchmark

To evaluate Metalink’s read-level taxonomic classification, we compared its detection sensitivity against MetaPhlAn4, which represents the most widely adopted marker-based approach. MetaPhlAn4 relies on clade-specific marker genes for taxonomic quantification, and its evaluation was restricted to reads successfully aligned to these markers based on Bowtie2^69^. We first benchmarked MetaPhlAn4 using its default database. Metagenomic reads were simulated from GTDB R207 genomes using InSilicoSeq^70^ with uniform parameters at 100 million reads, with three error models corresponding to HiSeq, NextSeq, and NovaSeq platforms. MetaPhlAn4-assigned SGB identifiers were converted to GTDB R207 taxonomy using mpa_vJun23_CHOCOPhlAnSGB_202403_SGB2GTDB.tsv. Classification accuracy was assessed by comparing direct read annotations against the original taxonomy of source genomes downloaded from the GTDB website.

The Metalink taxonomic database was constructed by integrating GTDB R220 prokaryotic and ICTV MSL38 viral genomes under a unified taxonomy framework combining GTDB R226 and ICTV MSL40. For benchmarking known species, Metalink was evaluated on prokaryotic genomes common to GTDB R207, R220, and R226, together with viral genomes common to ICTV MSL38 and MSL40, ensuring that these genomes were represented in the Metalink reference database (R220/MSL38), and maximizing their coverage by the MetaPhlAn4 marker database (with SGBs mapped to GTDB R207). For benchmarking novel species, Metalink was evaluated on prokaryotic genomes unique to R226 relative to both R220 and R207, and viral genomes unique to MSL40 relative to MSL38. Classification accuracy was assessed by comparing direct read annotations against the reference taxonomy of source genomes. For direct comparison, MetaPhlAn4 was benchmarked on the same known and unknown genome sets, although some of these genomes may already be represented in its built-in database.

The taxonomic binning accuracy was assessed using a read-level framework. For each read *i*, let *P_i_* denote the predicted taxonomic bin and *T_i_* the ground-truth taxonomic bin at the evaluated rank. Read-level true positives (*TP_i_*), false positives (*FP_i_*), and false negatives (*FN_i_*) were computed as the indicators of correct classification, spurious assignment, and missed true assignment, respectively. Global purity (precision) and F1 score were then computed by summing the *TP_i_*, *FP_i_*, *FN*_i_ across all reads:

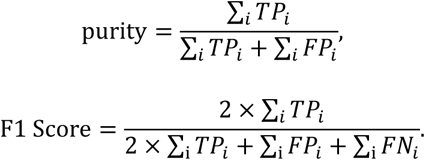

### Construction of hierarchical functional classification systems

We established two complementary functional hierarchies: LCFH, which leverages multi-KO co-assignment to infer parent-child relationships via the LCF principle, and SCFH, which constructs levels by hierarchical sequence clustering with cumulative frequency filtering. Both hierarchies use KOs as the minimal functional units at the lowest level. All genomes in the taxonomic database were subjected to gene prediction using Prodigal and Prodigal-GV^49^. Protein sequences with length greater than or equal to 33 amino acids were selected and annotated using EggNOG v5.0.2^71^ with DIAMOND v2.0.14.152^17^ (blastp mode, coverage >30% and *e* <1×10^-10^).

LCFH was constructed by reorganizing sequence KO assignment information of 45,231 unique KO annotations into a functional classification system. Multi-KO annotations serve as the names of parent nodes in the hierarchical tree structure, with parent-child relationships established by the LCF principle. This data-driven approach captures empirical co-annotation patterns while avoiding the combinatorial explosion of arbitrary KO permutations.

Formally, Let *K* be the set of all KO annotations. An LCFH node is a non-empty subset *n* ⊆ *K*. The initial level of node *n* is defined by its KO count:

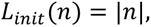

The inclusion relation ⊆ defines a partial order on the set of all observed nodes, denoted *N*_obs_. To convert this poset into a tree-structured hierarchy, we define the representation of node *n* at level *ℓ*, denoted *rep*(*n*, *ℓ*), as the identity of *n* after applying the unique parent rule at level *ℓ*. Let *N_ℓ_* denote the set of nodes at level *ℓ*, and let *C*(*n*, *ℓ*) be the set of candidate parent nodes at level *ℓ* that strictly contain *n*:

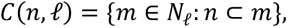

where m represents a candidate parent node. The node *n* is assigned to level *ℓ* according to the unique parent rule:

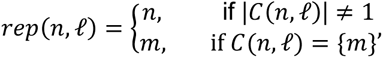

where ∣*C*(*n*, *ℓ*)∣ denotes the cardinality of the candidate set. This rule ensures that each node is assigned to a parent only when a unique containing node exists; otherwise, the node retains its identity at the current level (self-inheritance). The procedure is repeated iteratively from the initial level *L*_init_(*n*) upward to level 7.

LCFH construction proceeds in four steps. First, all KO annotations are collected and assigned to initial levels based on KO number. For example, lv3 contains multi-KO annotations with three KO members. Second, parent-child links are established by examining whether a node is a subset of nodes at the next higher level. Specifically, for each node at a given level, if a single node at the next higher level contains all its KO members, a link is constructed; otherwise the search continues upward level by level until a unique containing node is found or the root is reached. Third, this procedure is repeated for all nodes from lv2 to lvN-1. A critical constraint is that each child node must have exactly one parent node, ensuring that the classification system resolves KO hierarchical associations through LCF principle. Additionally, multi-KO annotations serve as node names for intermediate levels, while the sequences bearing these multi-KO annotations are also retained as leaf nodes at lv1 to enhance classification of ambiguous assignments. Fourth, missing node names at various levels are filled using the nearest lower-level node information, ensuring a complete tree-structured hierarchy.

SCFH was constructed by hierarchical clustering of protein sequences with different similarity thresholds. At each step, MMseqs2 v.13.45111^37^ was used to cluster sequences and generate representative sequences for the next lower-similarity threshold. For example, sequences clustered at 90% identity served as input for 70% clustering. Lower-similarity clusters were designated as parent nodes of higher-similarity clusters, yielding seven levels: KO_100%, KO_90%, KO_70%, KO_50%, KO_40%, KO_30%, KO_20%, with KO_90% and KO_50% corresponding to UniRef90 and UniRef50 clustering thresholds, respectively. At the KO_100% level, each cluster corresponds to one of the 45,231 unique KO annotations. Note that KO assignments of ORFs within low-level clusters (e.g., KO_100%) do not correspond to cluster boundaries at higher levels (e.g., KO_90%), because higher-level clusters may contain ORFs with diverse KO annotations. We therefore performed two-step filtering and aggregation to establish KO hierarchical relationships from sequence similarity clustering. First, representative KOs of ORFs within each cluster were selected based on cumulative frequency. For example, the KO set {KO1, KO2} from SCFH KO_100% level may correspond to multiple 100% similarity clusters. At the 90% level, these clusters may be grouped with other 100% clusters with distinct KO annotations (e.g., {KO1, KO2} or {KO1, KO3}), resulting in 90% clusters with heterogeneous KO compositions. The top 50% KOs by cumulative frequency within each 90% cluster were selected as the candidate node. Because multiple low-level clusters with identical KO sets may correspond to different upper-level clusters with distinct KO sets, the resulting KO hierarchy remains nested rather than tree-structured. Therefore, a representative parent node was then selected from multiple candidate parent nodes for each child node based on path-level cumulative frequency. For example, under a given node of KO_100%, the top 50% candidate parent nodes by cumulative frequency in KO_90% were selected as a KO set for the actual parent node. Hierarchies with filtering thresholds of 2%, 5%, and 10-100% at 10% intervals were tested. For simplicity, the same threshold was applied in both filtering steps. Thresholds exceeding 50% caused errors in downstream programs, and hierarchies with 2%-50% thresholds were therefore selected for subsequent testing.

Note that in LCFH, assignment to an intermediate level via LCF indicates that the sequence’s KO annotation is a subset of the named KO set of this level. In contrast, SCFH does not require unique parent node names, as each level represents a sequence set at a given similarity threshold rather than a name set. Consequently, SCFH assignment to an intermediate level indicates a representative set of KOs rather than complete functional equivalence or subset relationship to all constituent KOs of this level.

### Functional database construction, parameter sweeps and benchmark

Metalink functional databases were constructed using protein sequences from GTDB R226 prokaryotes, RefSeq Release 221 microeukaryotes, and ICTV MSL40 viruses. GTDB R220 genomes were used to construct benchmark databases with LCFH and SCFH as functional classification systems, respectively. Parameter sweeps were performed with varying *ℓ*, *k*, and *s* values. The lv1 level of LCFH was selected for database construction parameter optimization. The performance of LCFH and SCFH at different levels and the contribution of LCF to increased classified percent were subsequently evaluated.

### Functional binning benchmark

We used the intersection of GTDB R220 and R226 genomes to simulate known species and newly added genomes in GTDB R226 to simulate unknown species. Metagenomic reads were simulated using InSilicoSeq^70^ with uniform parameters at 100 million reads from nucleotide (ffn) sequences of both known and unknown species, with three error models corresponding to HiSeq, NextSeq, and NovaSeq platforms. KO annotations of eggNOG-Mapper for corresponding protein sequences served as the reference standard. We evaluated the accuracy of functional binning for Metalink, eggNOG-Mapper, and HUMAnN3 to fragmented metagenomic reads against the ground-truth KO annotations from complete ORF sequences. For each method, reads were processed as follows: Metalink performed read-level analysis on paired-end reads, where merged and unmerged reads were combined for ORF prediction and functional assignment via Kraken2 translated search against the custom LCFH protein database. Because Metalink leverages paired-end read information, which may provide additional phylogenetic signal for functional inference, we provided HUMAnN3 and eggNOG-Mapper with the identical read input (merged and unmerged paired-end reads) to ensure a fair comparison. EggNOG-Mapper with DIAMOND v2.0.14.152^17^ (blastp mode, coverage >30% and *e* <1×10^-10^) was run on the corresponding amino acid sequences of the same reads.

The functional binning accuracy was assessed using a sequence-level union-normalized framework. For each ORF *i*, let *E_i_* denote the set of predicted KOs and *K_i_* the set of ground-truth KOs from the corresponding complete ORF. True positives (*TP_i_*), false positives (*FP_i_*), and false negatives (*FN_i_*) were computed as the sizes of the intersection, set difference of prediction minus truth, and set difference of truth minus prediction, respectively. To ensure equal statistical weight regardless of the number of KOs assigned to an ORF, each ORF’s contribution was normalized by the size of the union of its predicted and true KO sets, ∣*E_i_* ∪ *K_i_*∣. The normalized per-ORF metrics were defined as:

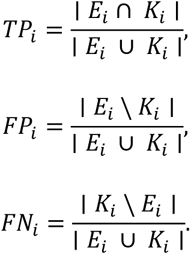

Global accuracy, purity (precision), completeness (recall), specificity, and F1 score were then computed by summing the normalized *TP_i_*, *FP_i_*, *FN*_i_, and *TN_i_* (true negatives, defined as the source ORF with empty eggNOG-Mapper annotation and empty read-level prediction) across all ORFs:

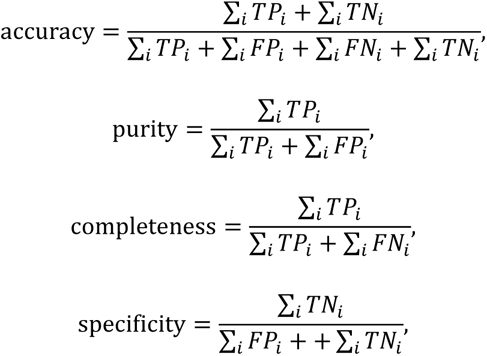

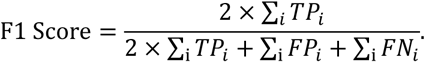

Additionally, to investigate performance across taxonomic and functional contexts, we stratified the evaluation by phylum and by KEGG metabolic pathway. For taxonomic stratification, ORFs were grouped according to their source genome’s taxonomic classification at the phylum rank, and the top 15 most abundant phyla were selected for comparison. For functional stratification, ORFs were grouped by the predominant pathway of their ground-truth KO annotations, with the top 15 most abundant pathways evaluated.

### Functional profiling benchmark

The CAMI II dataset^4^ was used for comparing profile-level accuracy across different tools. However, CAMI II was not designed for functional profiling benchmarking and lacked an appropriate gold standard. To address this limitation and enable reasonable comparison, we established a relative reference framework where different sequence types and tools were compared against each other. Specifically, the CAMI II metagenomes were divided into three groups based on sequence type and contig length thresholds: reads, contig 1000 bp, and contig 300 bp. Metalink and eggNOG-Mapper were then used to assign functional annotations to ORFs from reads, contig 1000 bp, and contig 300 bp, respectively. Bray-Curtis distances of KO-sample matrices were calculated using the R package vegan^72^. PCoA visualization was used to assess cross-tool and cross-sequence-type differences. Pairwise Bray-Curtis similarity for the same sample between tools and Pearson correlation of per-KO mean coverage were used to evaluate cross-tool consistency.

### Computational resource benchmarking

We benchmarked computational resource consumption for Metalink, eggNOG-Mapper, and HUMAnN3 in read-level functional binning using default parameters with 60 threads. Resource consumption for Metalink and RAT^29^ in mcr mode was also benchmarked on 10 CAMI II marine metagenome samples with 60 threads.

### Real-world datasets

The metagenomic dataset was assembled by searching the NCBI Sequence Read Archive (SRA) and the literature using the names of 29 target environments, retaining only paired-end shotgun metagenomes. Ten samples were selected per environment to ensure adequate representation. BioProjects individually contributing more than ten eligible samples were prioritized, and ten samples were randomly drawn from such projects. For environments where no single BioProject provided ten samples, the remaining samples were randomly supplemented from other BioProjects. This procedure yielded 290 samples from 29 environments across 70 BioProjects. The metatranscriptomic dataset comprised 108 seawater samples randomly selected from the TARA Oceans expedition^39,40^. For the Mantel test analysis, we retained the 95 samples with complete measurements across the 15 environmental factors examined.

### Other software and settings

Raw metagenomic and metatranscriptomic reads were trimmed using fastp v0.23.4^60^ to obtain clean reads. RiboDetector v0.3.1^73^ was used to remove rRNA sequences of metatranscriptomic reads. Default Metalink parameters were used in the benchmark unless otherwise specified. Metagenomic contigs of 300 bp and 1,000 bp were used to evaluate the performance of CAT_pack relative to Kraken2. For taxonomic benchmarking, the confidence threshold of Kraken2 was set to 0.6 for r mode, and 0.7 for c mode and for unmapped reads in integrated modes^36^. Quality of mMAGs was assessed using CheckM with the lineage_wf parameter and CheckM2, retaining genomes with ≥50% completeness and ≤1% contamination as taxonomic anchors for downstream multi-level integration. Only vMAGs classified as high-quality were retained.

## Acknowledgments

We thank Xinyue Wang, Lewen Liang, and Heng Wang for testing Metalink.

## Code availability

Metalink is available on GitHub (https://github.com/LiuyangLee/Metalink) and the Python Package Index (PyPI, https://pypi.org/project/metalink/). The custom R package mclink is available on Gith ub (https://github.com/LiuyangLee/mclink) and the Comprehensive R Archive Network (CRAN, https://cran.r-project.org/web/packages/mclink/index.html).

## Author Contributions

L.L. designed the research, developed the software code, performed the analyses and benchmarking, and wrote the paper. Y.W. designed the research, critically revised the paper and provided funding.

---

**Extended Figure 1.**
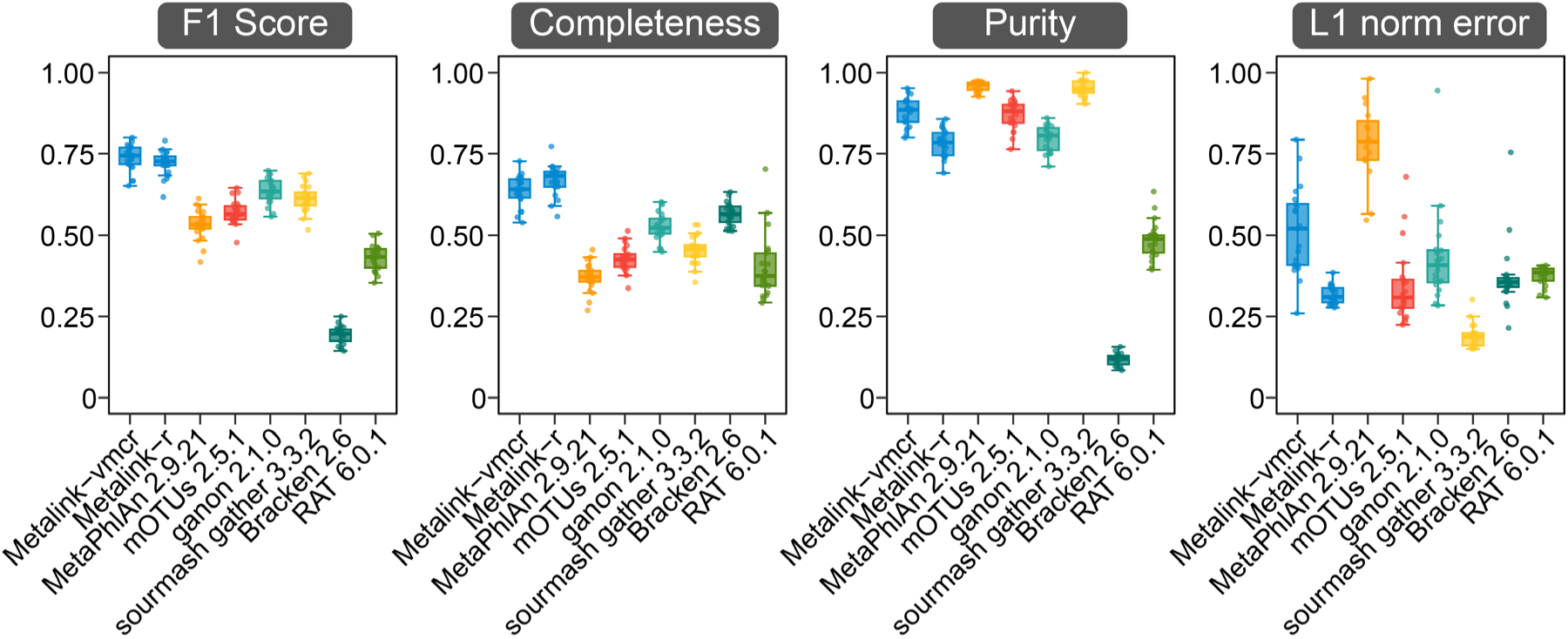
Taxonomic profiling performance of Metalink-vmcr and Metalink-r modes on CAMI II plant-associated datasets at the genus level (n = 21 samples).

**Extended Figure 2.**
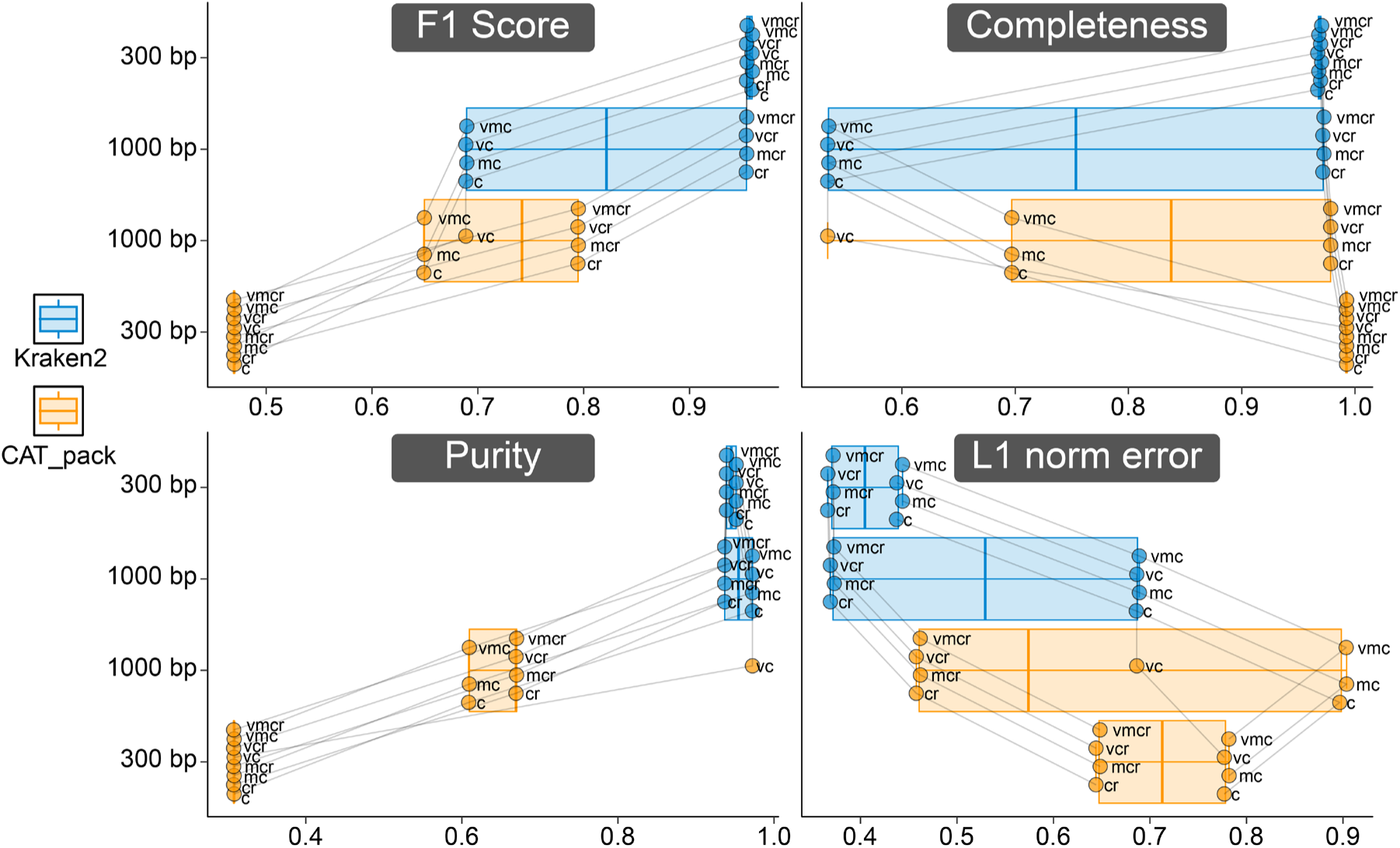
Impact of contig taxonomy classification method and length threshold on taxonomic profiling performance. Comparison of different taxonomic classification approaches (Kraken2 vs. CAT_pack) for contigs and evaluation of length thresholds (1,000 bp vs. 300 bp) to taxonomic profiling performance in Metalink.

**Extended Figure 3.**
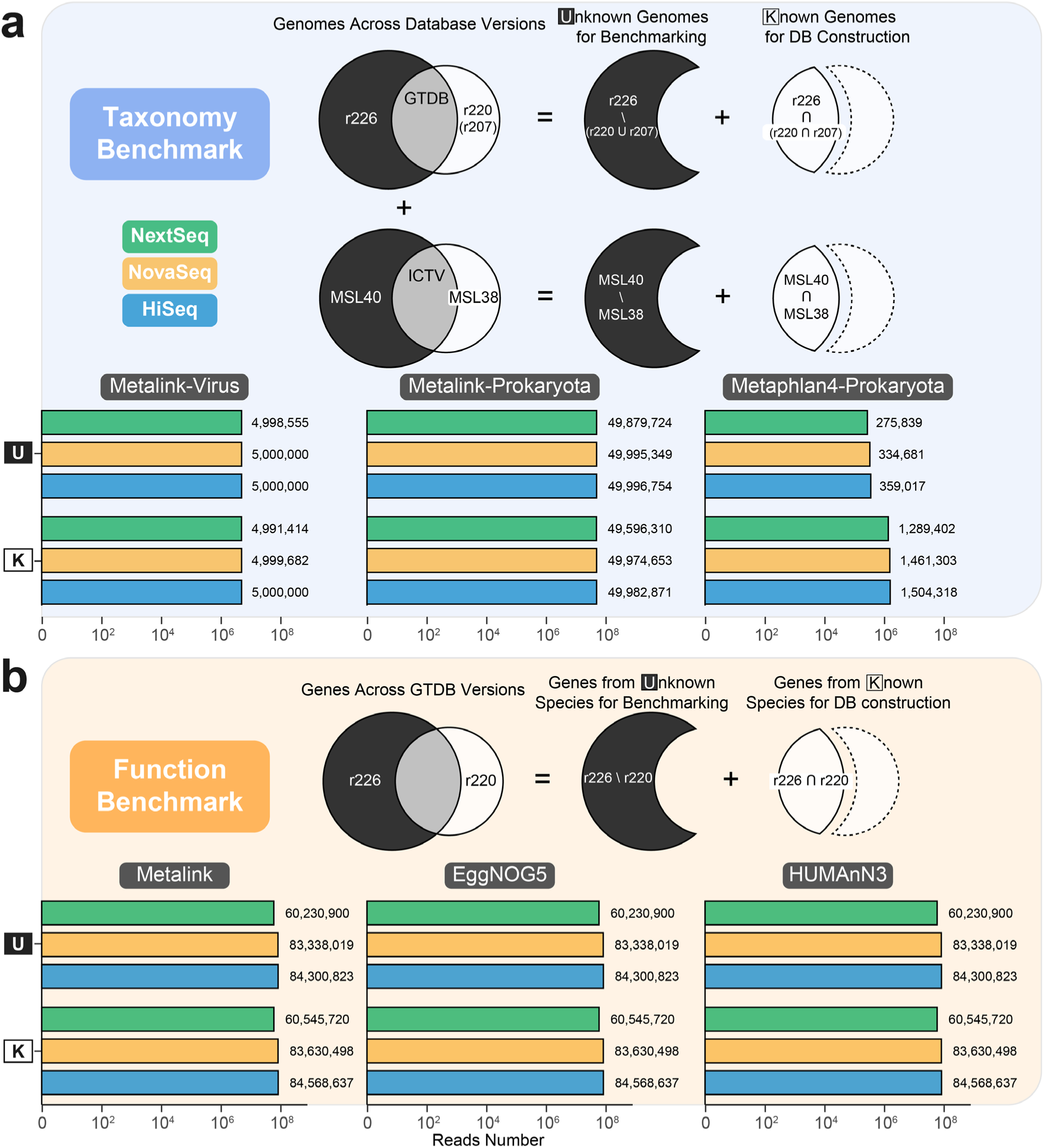
Benchmarking of taxonomic and functional binning using simulated read datasets. a, Taxonomic benchmarking across three sequencing platforms (HiSeq, NovaSeq, and NextSeq). For Metalink-Virus and Metalink-Prokaryota, all simulated reads were used. For MetaPhlAn4, only reads aligning to its marker genes were retained after inputting all reads. Venn diagrams illustrate prokaryotic genome overlaps across GTDB versions (r207, r220, and r226) and viral genome overlaps across ICTV versions (MSL38 and MSL40). The Metalink database for benchmarking was constructed using prokaryotic genomes from GTDB r220 and viral genomes from ICTV MSL38, classified under the integrated GTDB r226 and ICTV MSL40 taxonomy. Unknown genomes used for benchmarking novel species are defined as prokaryotic genomes present in r226 but absent from both r220 and r207, and viral genomes present in MSL40 but absent from MSL38. Known genomes used for benchmarking known species are defined as prokaryotic genomes present in the intersection of all three versions, and viral genomes present in both MSL38 and MSL40. **b,** The total number of simulated reads for functional benchmarking across Metalink, eggNOG-Mapper (EggNOG5), and HUMAnN3. Venn diagrams illustrate gene overlaps across GTDB versions (r220 and r226), distinguishing genes of known species from those of unknown species. Whereas the standard Metalink functional database is constructed from GTDB R226 genomes, a dedicated benchmark database was constructed from R220 genomes, so that genes of species newly added in R226 serve as the unknown test set; genes of species present in both versions (R220 ∩ R226) were used for benchmarking known functions.

**Extended Figure 4.**
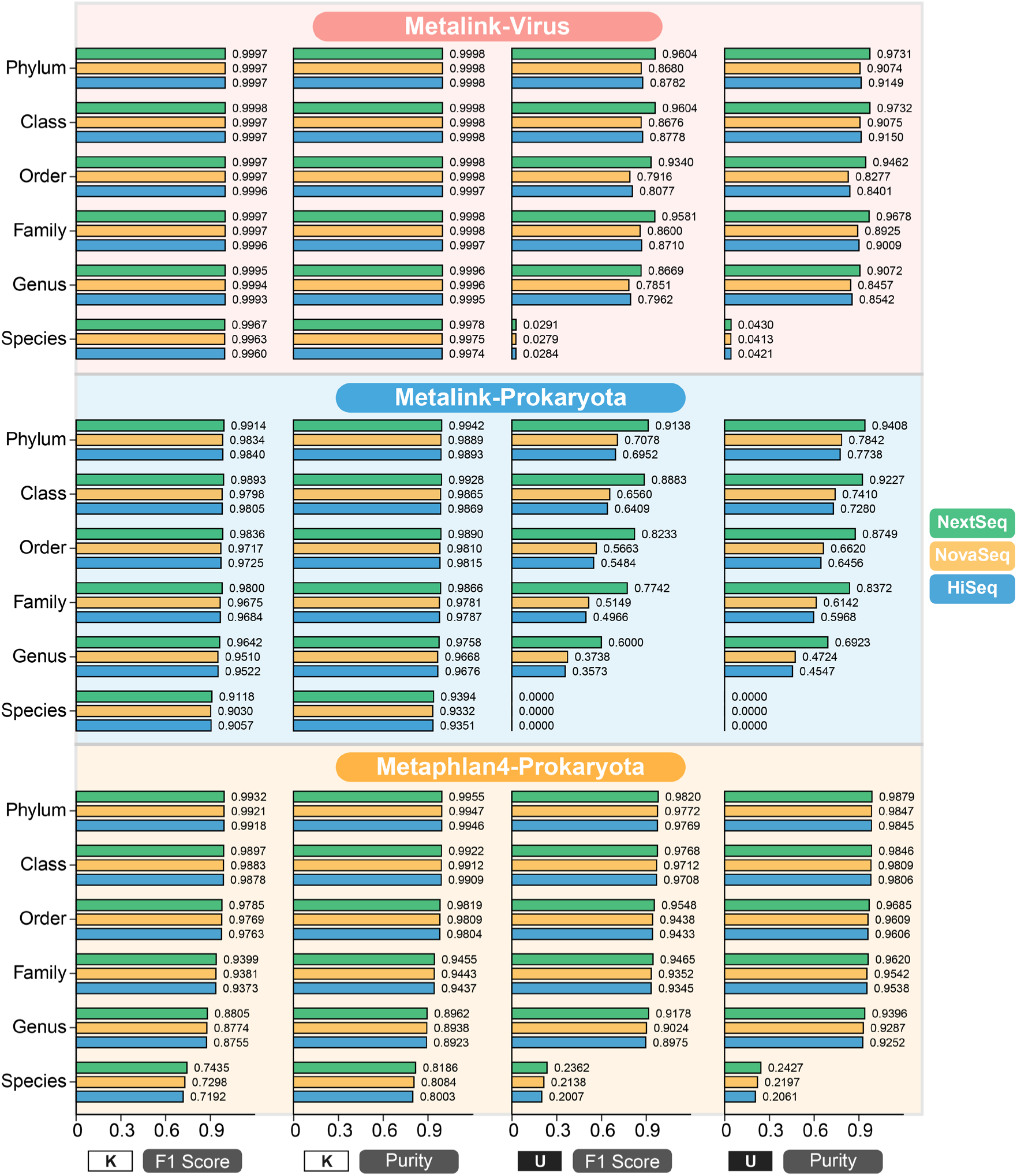
Benchmarking of taxonomic binning of Metalink-r using a split-database approach to distinguish known and unknown species. Note that the “unknown” designation may not be fully applicable to MetaPhlAn4: its marker set is organized around SGBs that do not correspond one-to-one to versioned GTDB genomes, so whether genomes in the “unknown” set are already represented in its built-in marker database cannot be assessed exactly. Performance metrics (F1 score and purity) are shown across taxonomic ranks from phylum to species for both known and unknown genome sets.

**Extended Figure 5.**
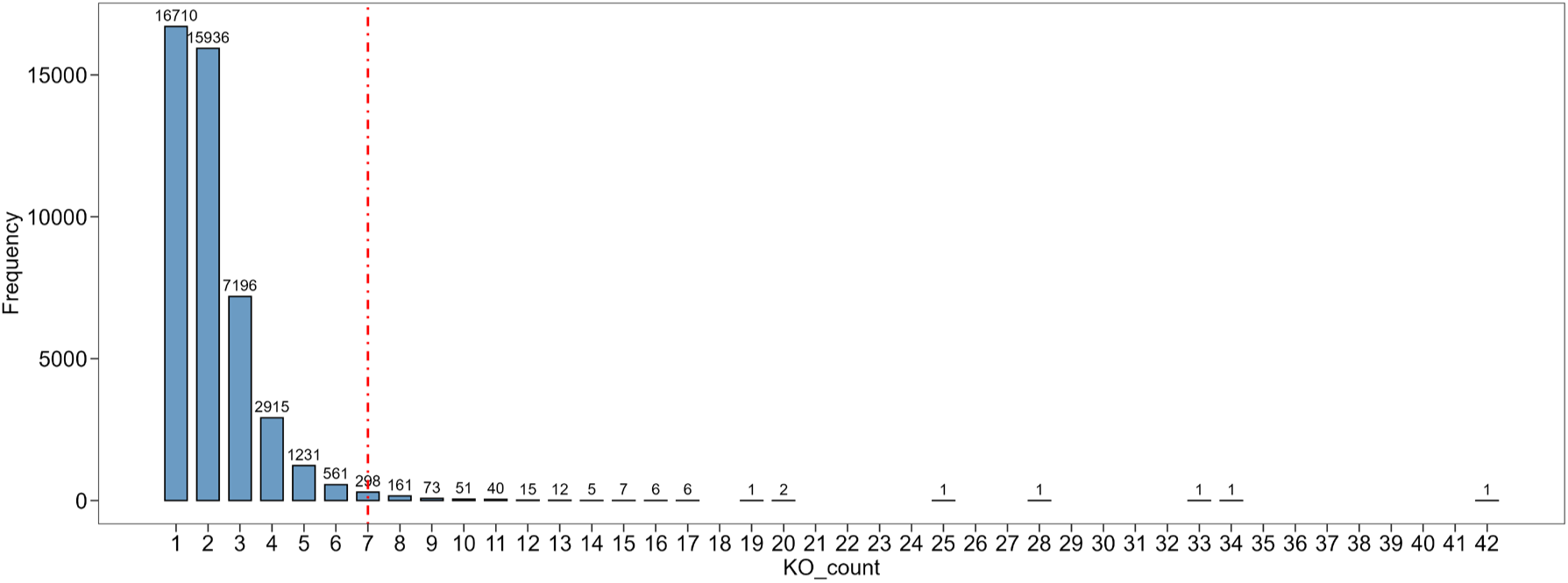
Frequency distribution of unique KO counts contained in nodes at the lv1/KO_100 level for the Last Common Function-based functional Hierarchy (LCFH) and the Similarity Clustering with Frequency-filtered functional Hierarchy (SCFH).

**Extended Figure 6.**
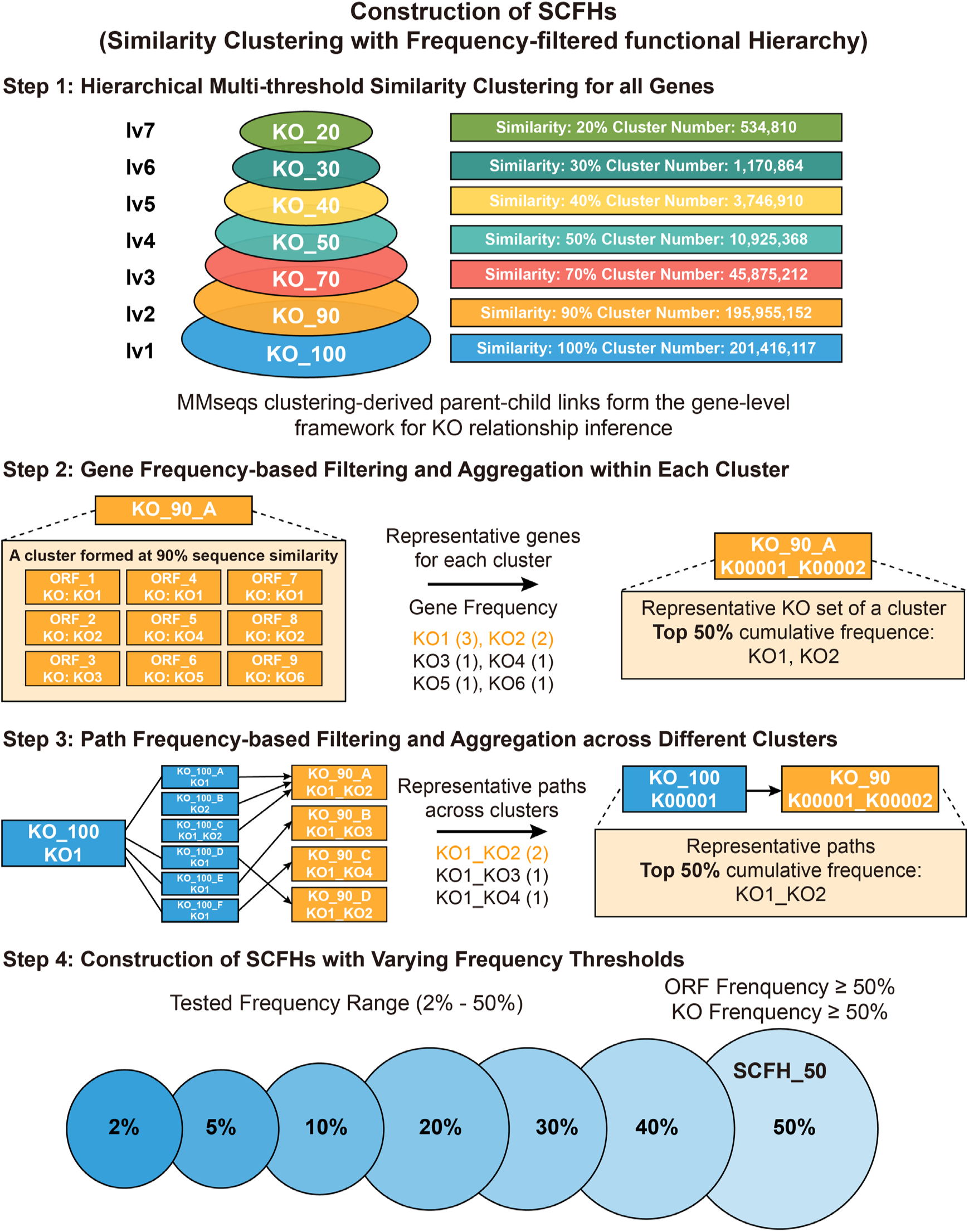
Overview of the Similarity Clustering with Frequency-filtered functional Hierarchy (SCFH).

**Extended Figure 7.**
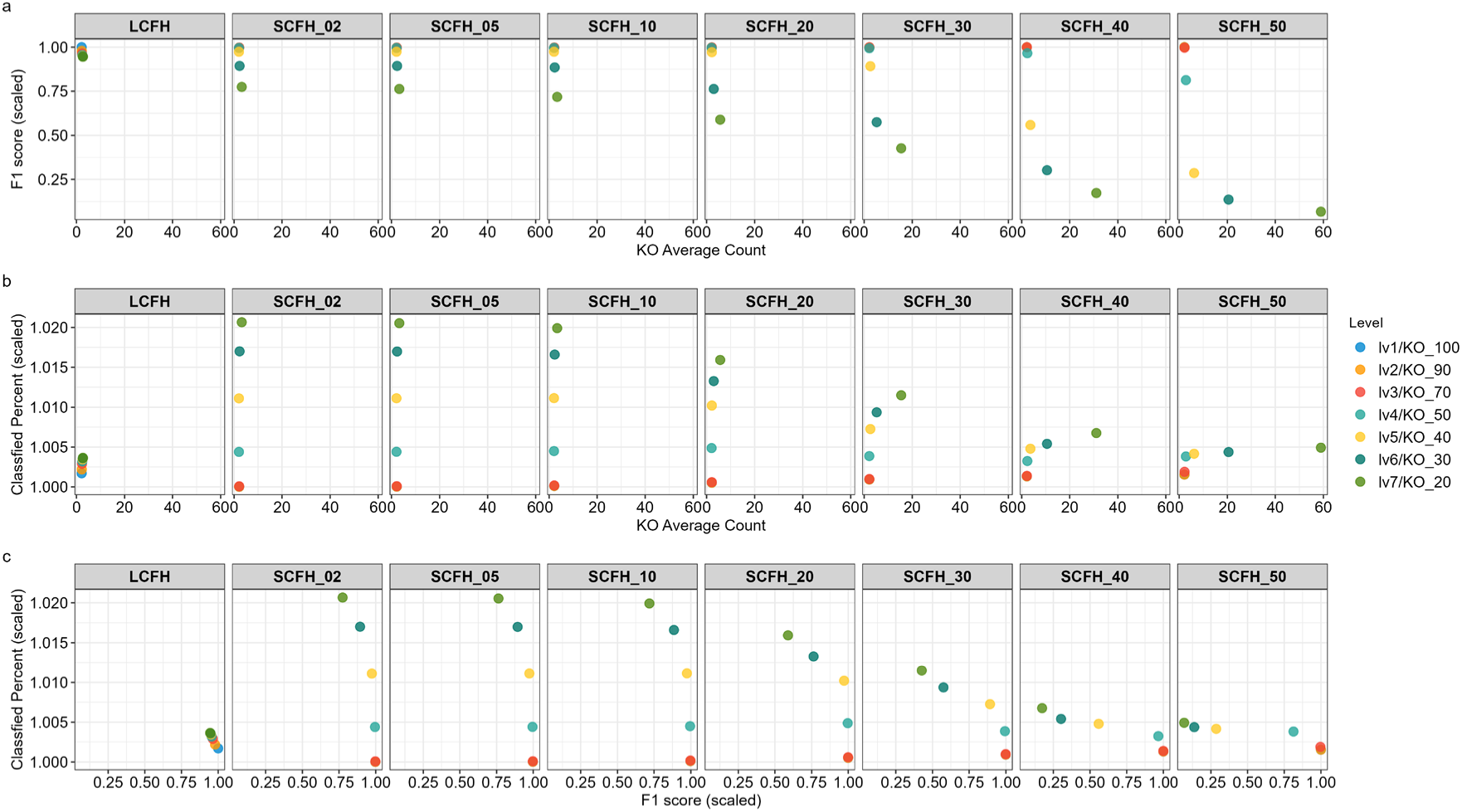
Benchmark of the Last Common Function-based functional Hierarchy (LCFH) and the Similarity Clustering with Frequency-filtered functional Hierarchy (SCFH), considering F1 score (per-read), classified percentage (overall), and node name complexity across different levels.

**Extended Figure 8.**
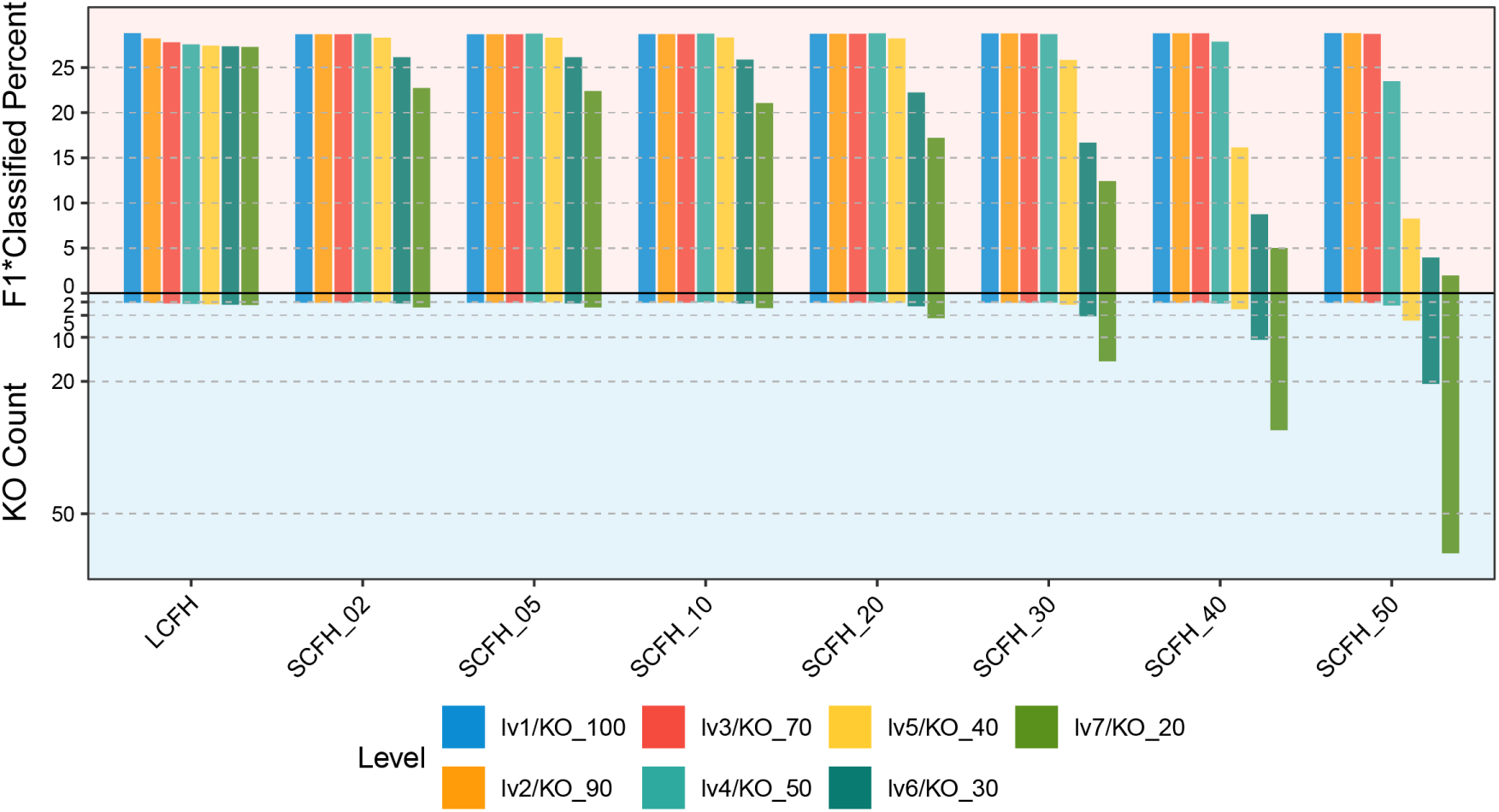
Benchmark of the Last Common Function-based functional Hierarchy (LCFH) and the Similarity Clustering with Frequency-filtered functional Hierarchy (SCFH), using the product of F1 score and Classified Percent as the integrated metric, along with node name complexity across different levels.

**Extended Figure 9.**
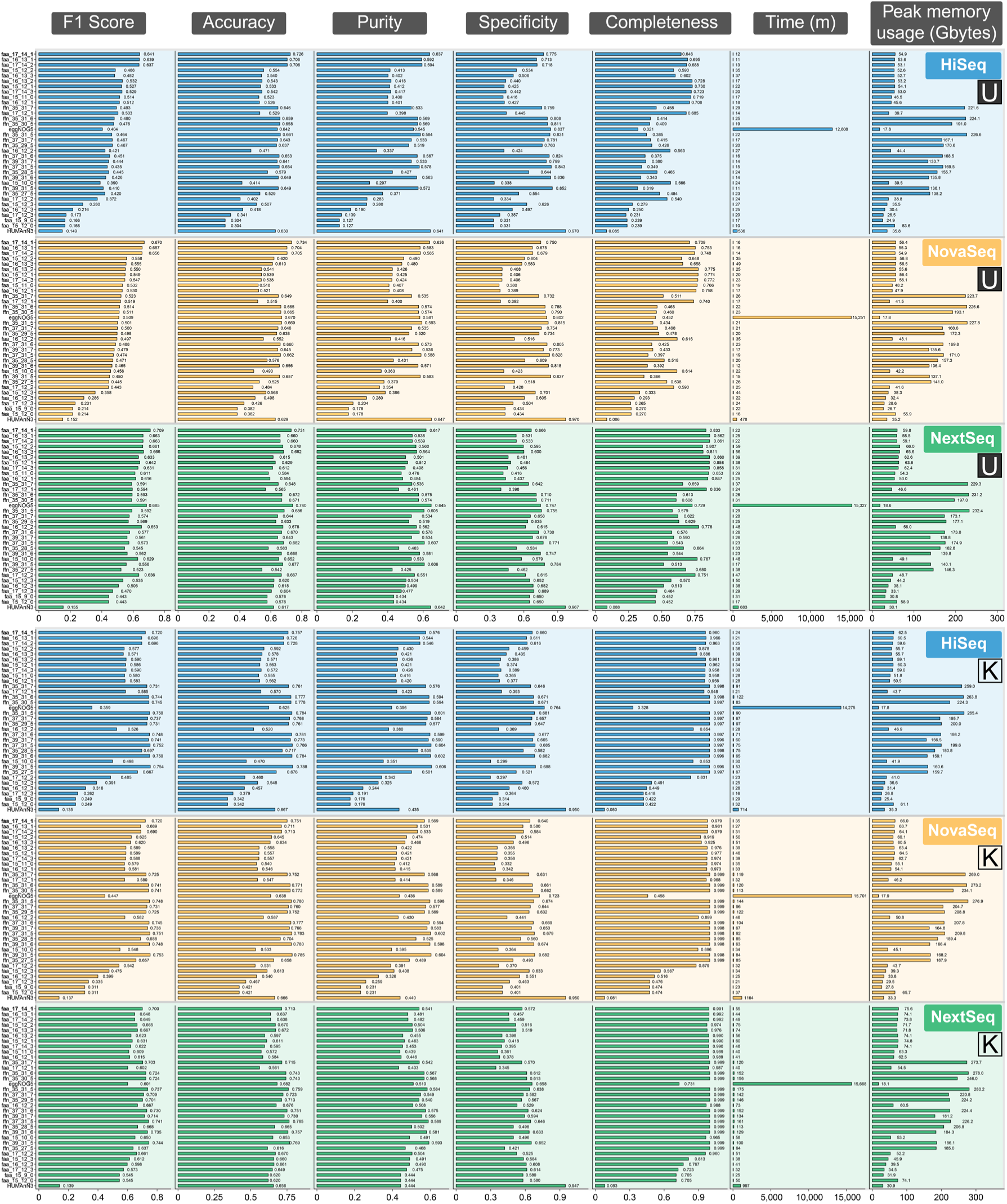
Parameter optimization for functional binning database construction using Metalink-r. U, unknown species; K, known species. Systematic scanning of parameters (*ℓ*, *k*, and *s*) was performed for both nucleotide (ffn) and protein (faa) databases. Genomes from known (older GTDB version) and unknown (newer GTDB version) species were fragmented using error models simulating three sequencing platforms (HiSeq, NovaSeq, NextSeq). Benchmarking was conducted against eggNOG-Mapper and HUMAnN3.

**Extended Figure 10.**
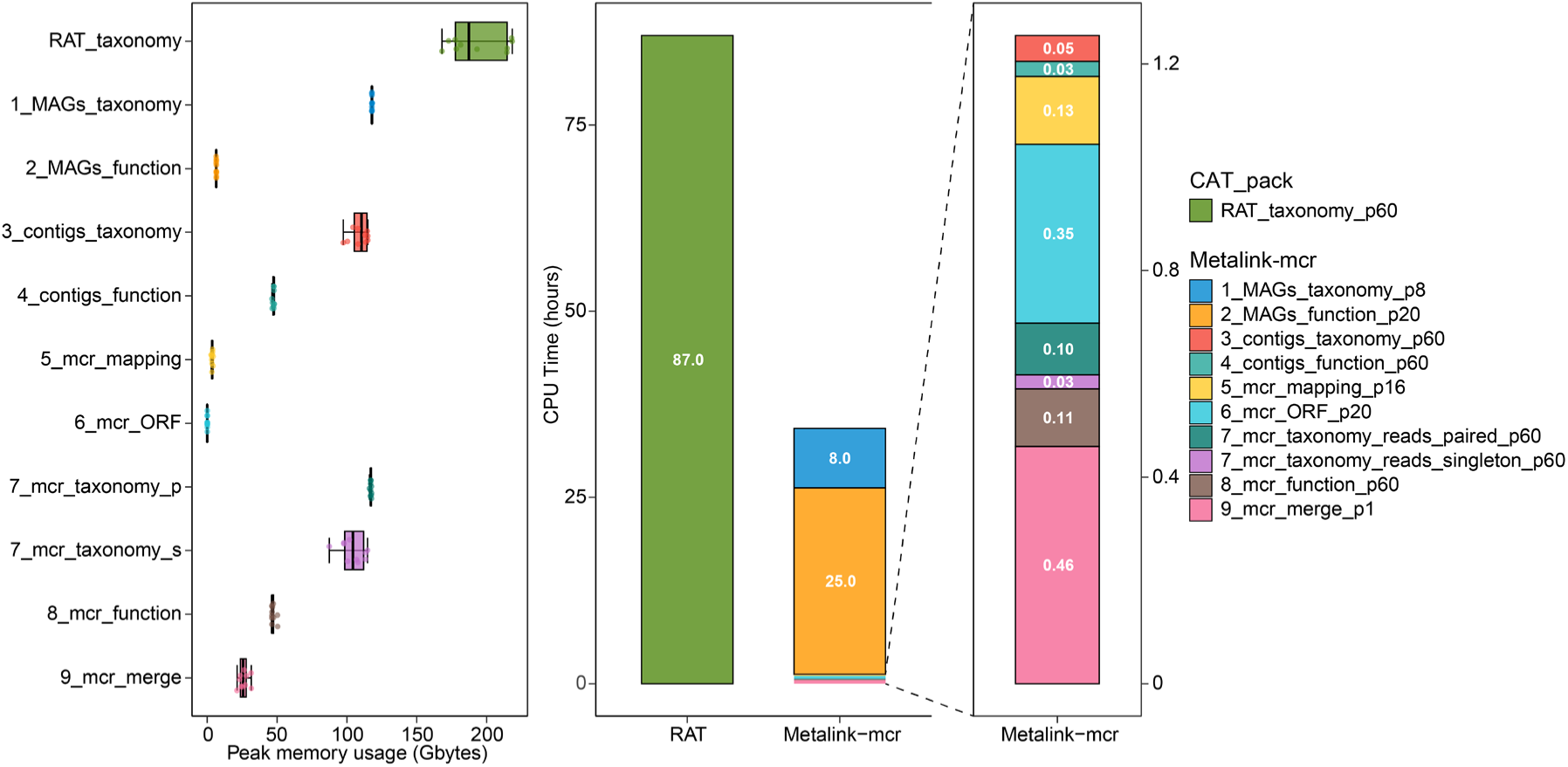
Computational resource benchmarking. Peak memory usage and CPU time consumption across nine analytical steps of Metalink-mcr compared to total metrics of RAT (CAT_pack reads --mode mcr). Metalink-mcr was configured with a maximum of 60 threads, with individual steps optimized to use fewer threads; RAT was executed with 60 threads. All measurements were obtained from 10 CAMI II marine metagenome samples. Metalink-mcr performs integrated taxonomic and functional analysis, whereas RAT provides taxonomic annotation only. The nine steps of Metalink-mcr are: (1) MAG taxonomic annotation with BAT, (2) MAG functional annotation with eggNOG-Mapper, (3) contig taxonomy classification with Kraken2, (4) contig functional annotation with Kraken2, (5) read mapping with BWA-MEM2, (6) open reading frame (ORF) prediction with FragGeneScanRs and prodigal, (7) unmapped read taxonomy classification (paired-end and singleton reads), (8) unmapped read functional annotation, and (9) final taxonomy-function integration. Peak memory usage is displayed as boxplots with individual data points (jittered). CPU time is shown as stacked bar plots, with each segment representing the contribution of individual Metalink-mcr steps or total RAT runtime.

