## Supplemental Figures 1 to 6 for "Metalink: Joint taxonomic and functional profiling of metagenomes and metatranscriptomes across resolution scales"

*Yinzhao Wang

**This file includes:**

Figures S1 to S6

**Other supporting materials for this manuscript include the following:**

Supplementary Data 1 and 2

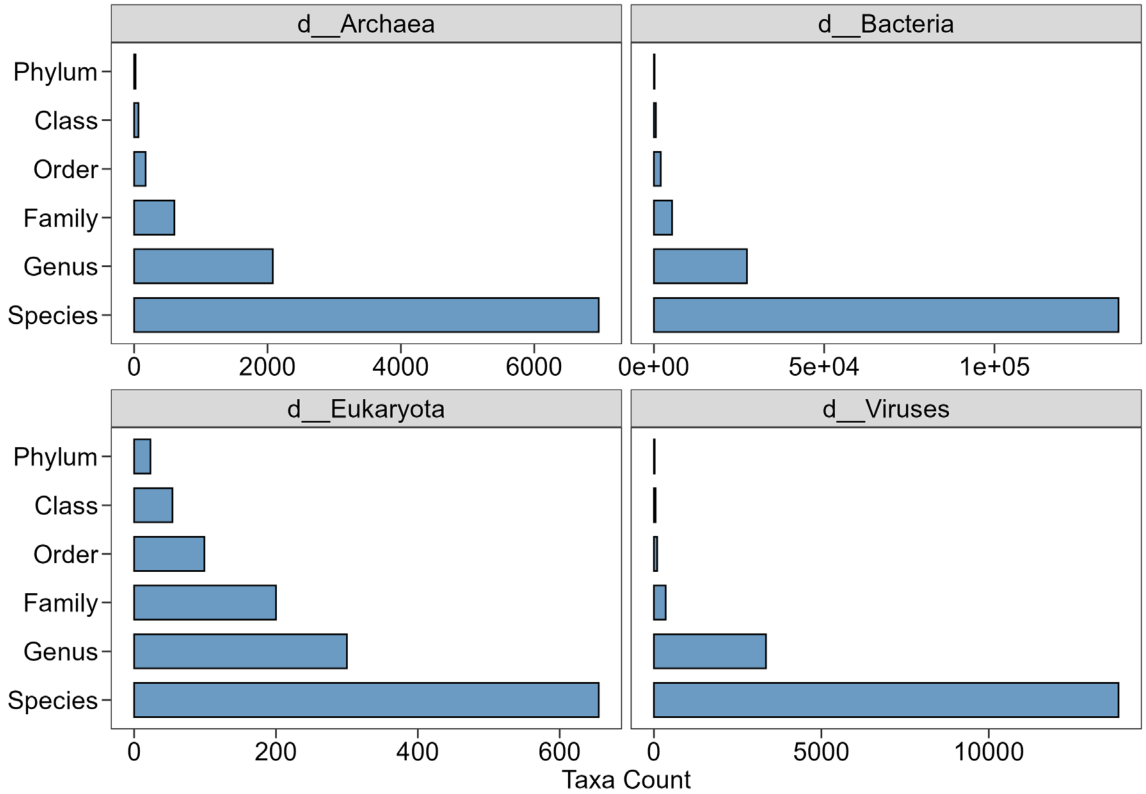


**Figure S1.** Number of unique nodes from phylum to species level in the integrated taxonomic hierarchy.

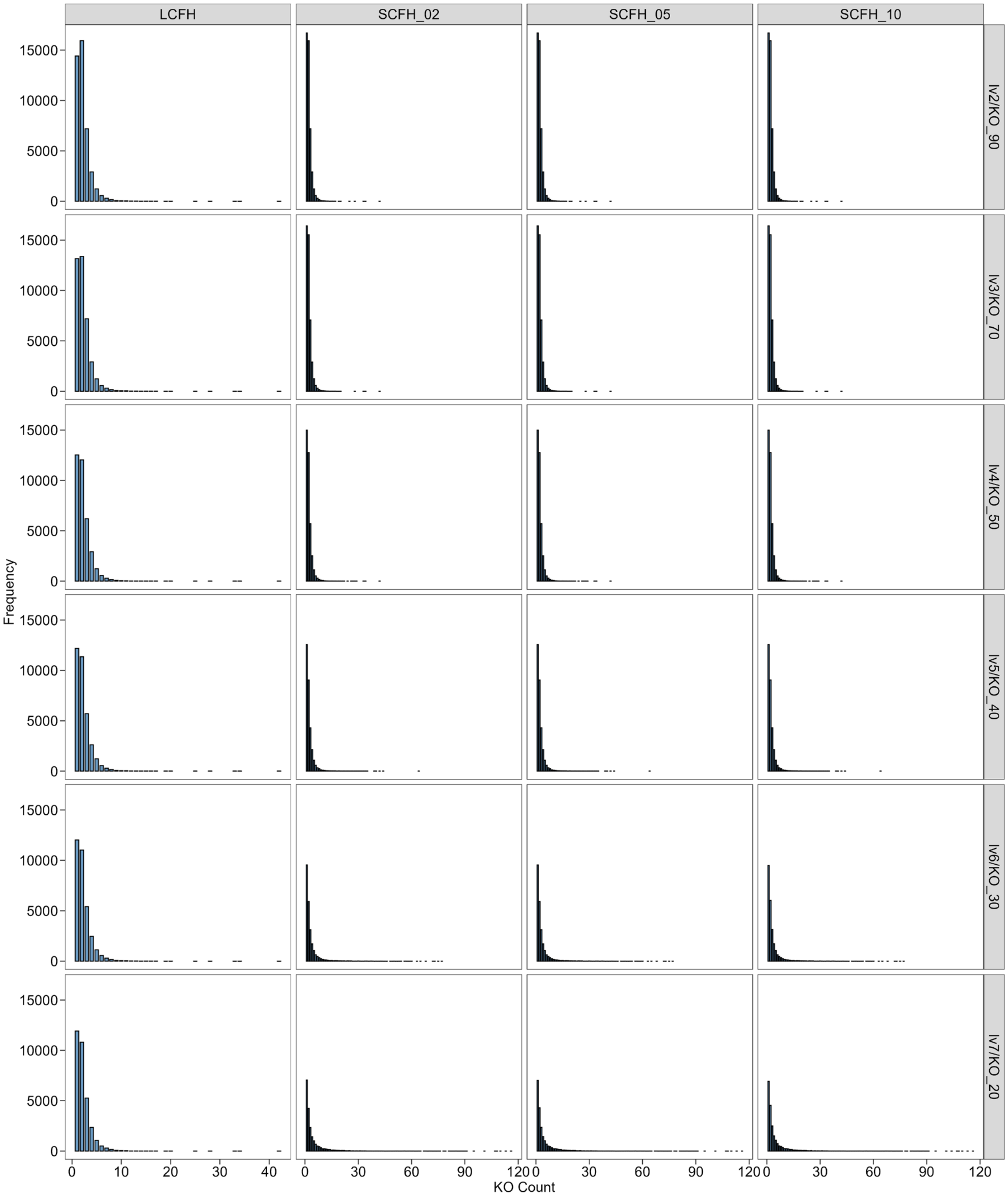


**Figure S2.** Frequency distribution of KO counts across all nodes at each level from lv2/KO_90 to lv7/KO_20 for the Similarity Clustering with Frequency-filtered functional Hierarchy (SCFH) at 2%, 5%, and 10% thresholds, and the Last Common Function-based functional Hierarchy (LCFH).

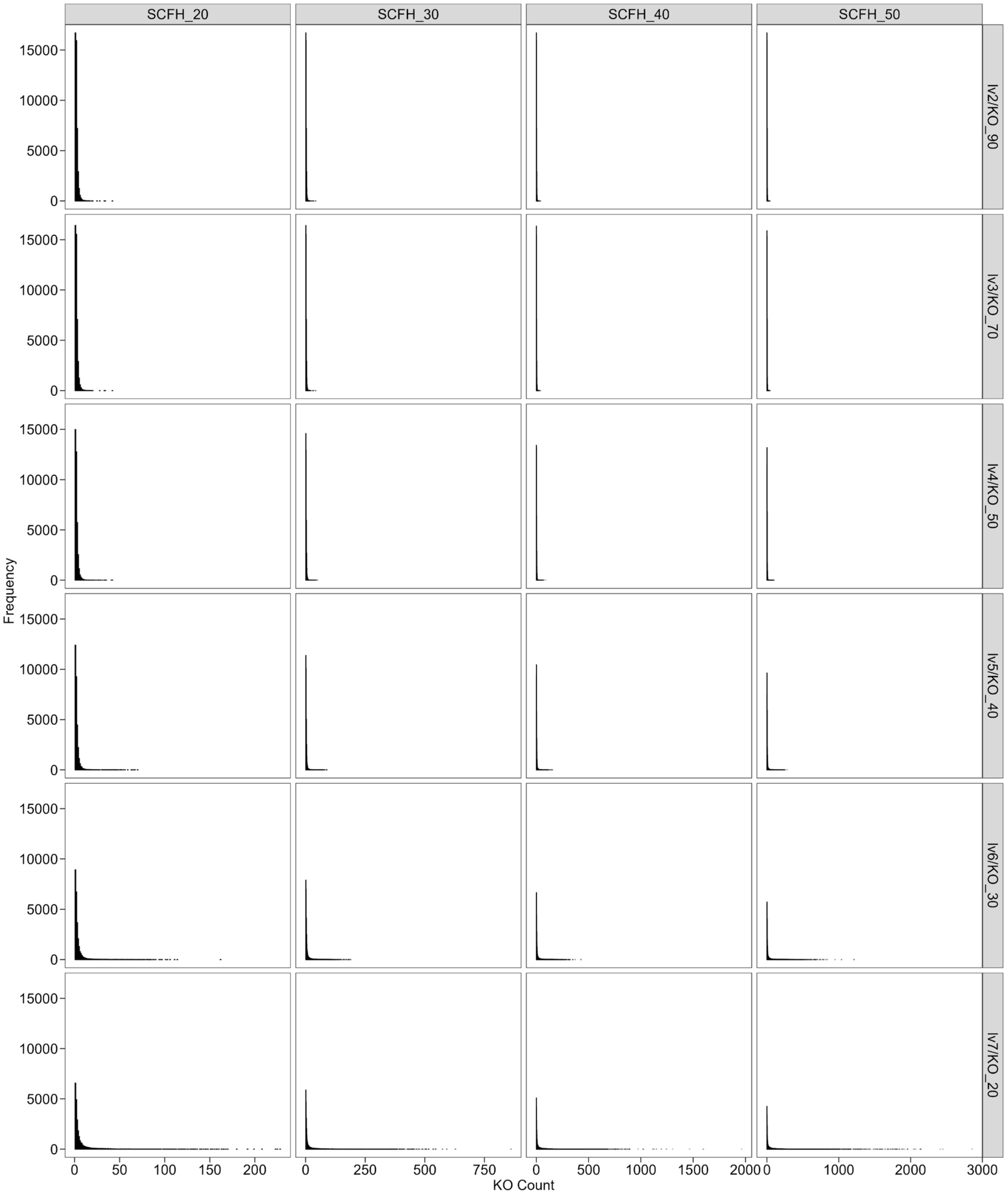


**Figure S3.** Frequency distribution of KO counts across all nodes at each level from lv2/KO_90 to lv7/KO_20 for the Similarity Clustering with Frequency-filtered functional Hierarchy (SCFH) at 20%, 30%, 40%, and 50% thresholds.

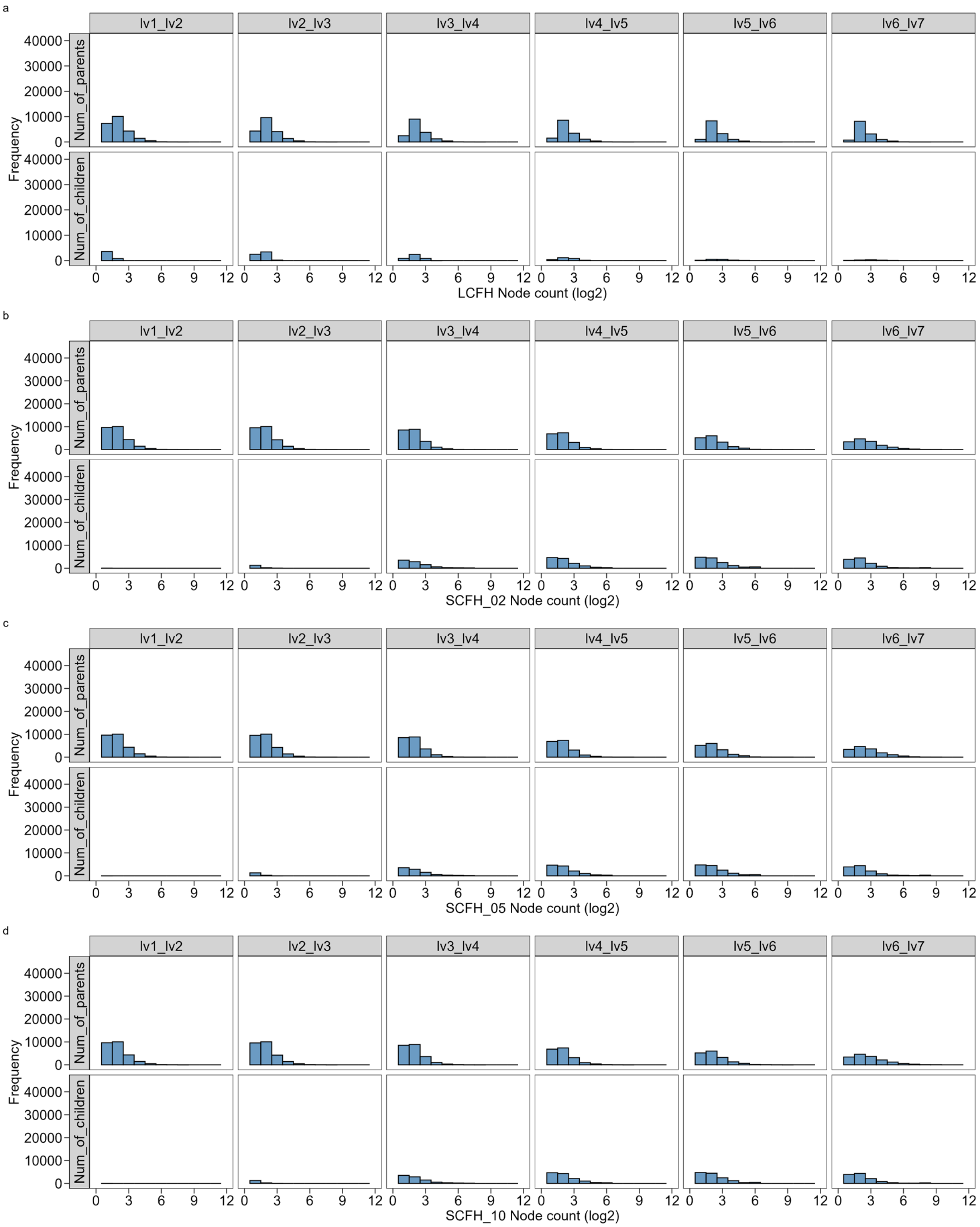


**Figure S4. Structural properties of hierarchical functional classification systems.** The number of unique parent nodes sharing identical KO content with distinct lower-level nodes (Num_of_parents, upper row) and the number of unique paths connecting parent and child nodes (Num_of_children, lower row) between adjacent levels of the Similarity Clustering with Frequency-filtered functional Hierarchy (SCFH) at 2%, 5%, and 10% thresholds, and the Last Common Function-based functional Hierarchy (LCFH).

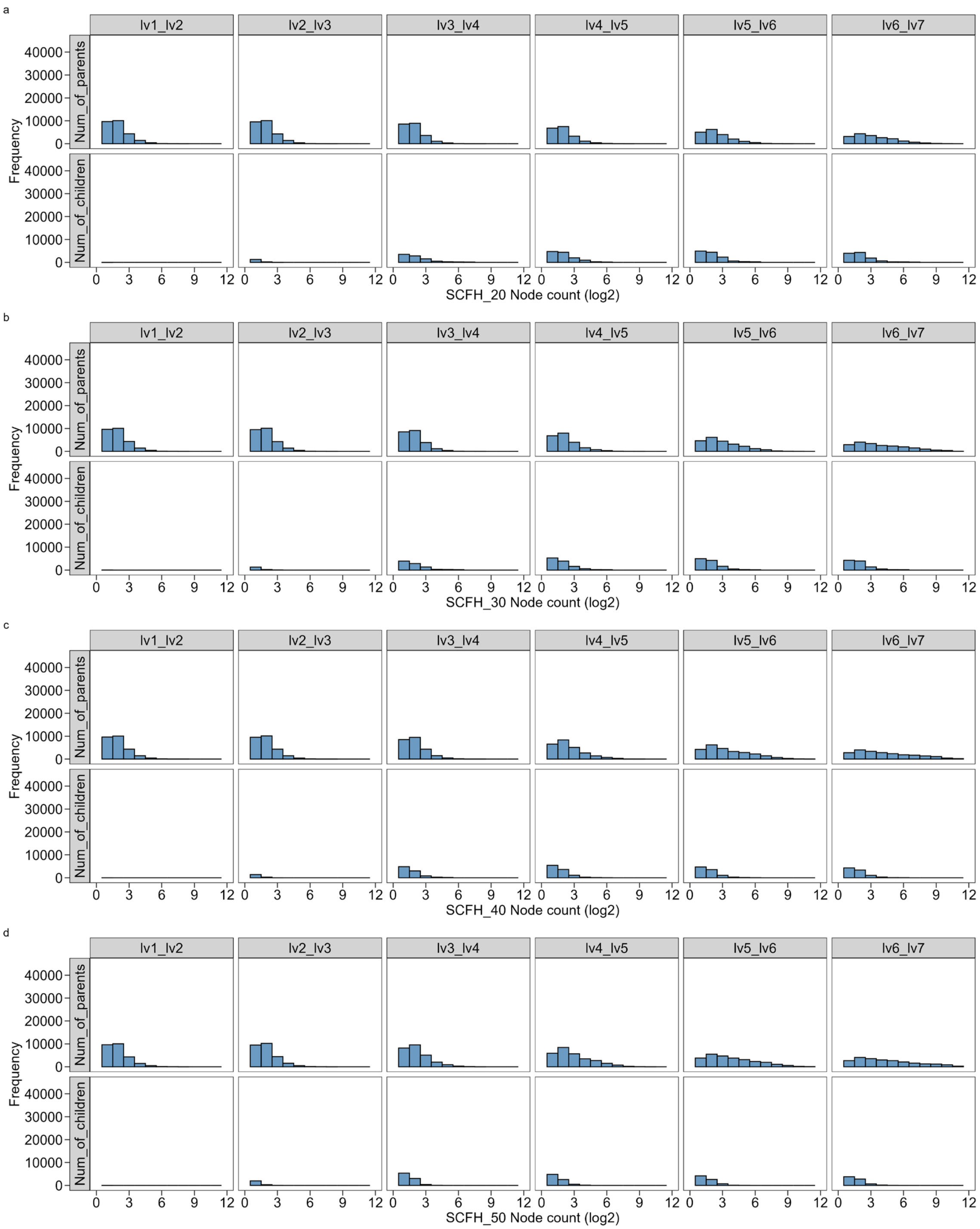


**Figure S5. Structural properties of hierarchical functional classification systems.** The number of unique parent nodes sharing identical KO content with distinct lower-level nodes (Num_of_parents, upper row) and the number of unique paths connecting parent and child nodes (Num_of_children, lower row) between adjacent levels of the Similarity Clustering with Frequency-filtered functional Hierarchy (SCFH) at 20%, 30%, 40%, and 50% thresholds.

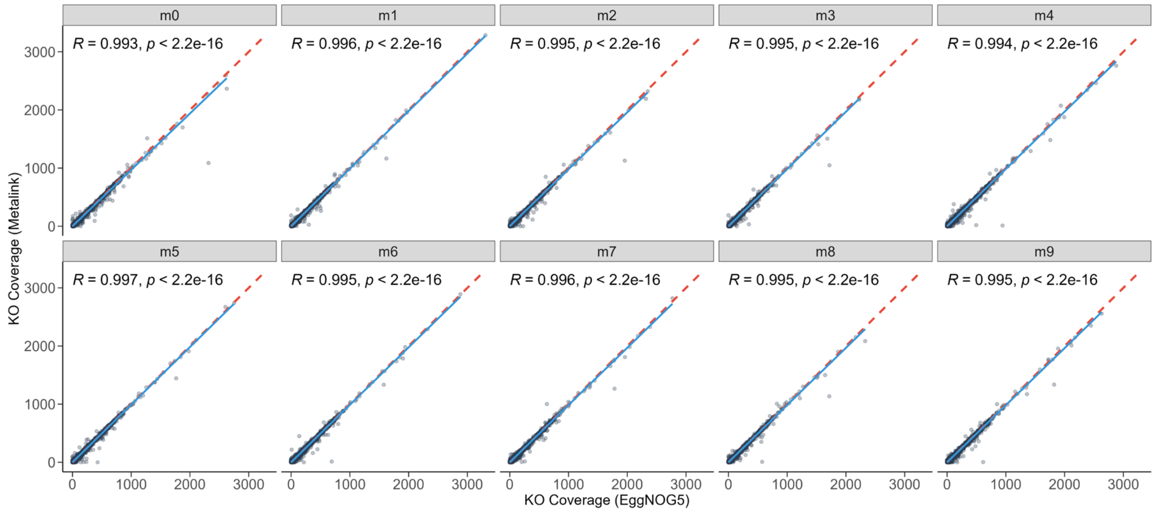


**Figure S6. Consistency between Metalink and eggNOG-Mapper (EggNOG5) annotations for CAMI II marine metagenomic samples at the contig 1,000 bp level.** Each panel shows a pairwise comparison between two software tools. Each point represents a KO, with the x- and y-axes showing its mean coverage annotated by the respective tools.
